# A molecular dynamics investigation of CDK8/CycC and ligand binding: conformational flexibility and implication in drug discovery

**DOI:** 10.1101/169581

**Authors:** Wei Chen, Zhiye Tang, Tim Cholko, Chia-en A. Chang

**Affiliations:** Department of Chemistry, University of California, Riverside, CA92521

**Keywords:** Cyclin-dependent kinase 8, Cyclin C, Molecular Dynamics, DMG, ligand binding, protein motions

## Abstract

The activities of CDK8 with partner Cyclin C (CycC) are a common feature of many diseases, especially cancers. Here we report the study of dynamic behaviors and energy profiles of 13 CDK8/CycC systems, including the DMG-in and DMG-out conformations as well as 5 type I ligands and 5 type II ligands, with all-atom unbiased molecular dynamics (MD) simulations. We observed numerous regional motions within CDK8, which move in concert to form five major protein motions. The motion of the activation loop doesn’t appear to influence the binding of both types of ligands. Type I ligands remarkably reduce the motion of the C-terminal tail through the strong cation-π interaction between the ligands and ARG356, and type II ligands stabilize the αC helix by forming stable hydrogen bonds with GLU66. The MD calculations also confirmed the importance of CycC to the stability of the CDK8 system as well as the ligand binding. The MMPB/SA results show that van der Waals interaction is the main driving force for the binding of both types of ligands, but electrostatic energy and entropy penalty plays important roles in the binding of type II ligands. The volume analysis results indicate that the induced fitting theory applies in the binding of type I ligands. These results would help to improve the affinities of the existing ligands. Our MD work is complementary to crystal structures and may have implications in the development of new CDK8 inhibitors as well as in the field of drug discovery.

## 1. Introduction

Cyclin-dependent kinases (CDKs) are among the major regulators of cell cycle and transcription [1]. The functions of CDKs depend on the binding with regulatory proteins called cyclins. CDK8 together with Cyclin C (CycC), MED12 and MED13 forms a regulatory kinase module of the mediator complex [2–4], which is a large protein assembly that couples gene-specific transcriptional regulators to the general RNA polymerase II transcription machinery [5, 6]. A number of studies have showed that CDK8 modulates the transcriptional output from distinct transcription factors involved in oncogenic control [7]. These factors include the wnt/β-catenin pathway, Notch, p53, and TGF-β [8, 9].

Due to the fundamental biological roles that CDKs play, it is not surprising that their activities are a common feature of many diseases, especially cancers. CDK8 has recently attracted considerable attention in recognition of its key roles in oncogenesis. Gene expression of CDK8 is related with the activation of β-catenin which is a core transcriptional regulator of canonical wnt signaling in gastric cancers [10–12]. CDK8 has been identified to be essential in cell proliferation in melanoma and to act as an oncogene in colon cancer as the CDK8 gene is amplified in about 60% of colorectal cancers [13, 14]. CDK8 gene expression also correlates with prognosis in breast and ovarian cancers [15]. Additional cancer-relevant activities of CDK8 include growth factor-induced transcription [16], modulation of TGF b signaling [17] and phosphorylation of the Notch intracellular domain [18, 19].

The research on CDK8 ligands is still on its early stage, as evidenced by the lack of clinical trial data [20]. The steroidal natural product cortistatin A was the first-reported high affinity and selective ligand for CDK8 with an IC50 value of 12 nM in vitro and the complete selectivity against 387 kinases [21]. The existing ligands fall into two categories based on the major conformations of CDK8 they bind to. Type I ligands bind to DMG-in (Aspartate-Methionine-Glycine near the N-terminal region of the activation loop) conformation and occupy the ATP-binding site. The Senexin-type, the newer CCT series and COT series compounds, which possess 4-aminoquinazoline [22], 3,4,5-trisubstituted pyridine [23] and 6-azabenzothiophene [24] scaffolds, respectively, belong to this category. Type II ligands bind to DMG-out conformation and occupy the ATP-binding site and the allosteric site (deep pocket). The deep pocket is adjacent to the ATP-binding site and accessible in CDK8 by the rearrangement of the DMG motif from the active (DMG-in) to the inactive state (DMG-out). This pocket is inaccessible in the active conformation (DMG-in), where the Met174 side-chain is reoriented to make the site available to ATP [20]. Typical type II CDK8 ligands are sorafenib and imatinib analogs that contain an aryl urea core [25]. The research on CDK8 ligands keeps highly active. Very recently 4,5-dihydrothieno[3’,4’:3,4]benzo[1,2-d] isothiazole derivatives were found to have sub nanomolar in-vitro potency (IC50: 0.46 nM) against CDK8 and high selectivity [26].

Since Scheneider et. al revealed the first crystal structure for human CDK8/CycC complexed with sorafenib (PDBID: 3RGF) in 2011 [27], up to date there are 24 crystal structures available for this kinase system, thus providing plenty of structural information for the computational approaches to understand the molecular functions and interactions with substrates and ligands of CDK8. When compared with other CDKs, CDK8 displays additional potential recognition surfaces for interactions, possibly for the recognition of MED12, MED13, or the substrates of CDK8. However, all of the crystal structures are lack of part or all of the residues within the activation loop which is critical to ATP substrate and ligand binding. The activation loop is quite flexible. Without the structural details of the activation loop, it is unclear how its motion affects other regions of the protein and its impact on the stability of the protein structure and ligand binding. Furthermore, CDK8 relies on its partner CycC to function. It would be of interest to find out the implications of CycC on the dynamics of the CDK8 regions, especially those near the binding sites. If the influence is negligible, CycC could be ignored and computations on CDK8 would be significantly sped up. Otherwise CycC must be considered in the system to keep the calculations meaningful. Such information is not available from the crystal structures as well. Moreover, although crystal structures provide the binding modes of ligands to proteins, the stability of the binding modes and the possibility of the alternative binding poses remain unknown. This knowledge is valuable for the improvement of the existing ligands. The computational methods such as Molecular Dynamics present complementary approaches to understand the details of structural changes in the course of ligand binding to proteins. Wu Xu, et. al presented 50 ns of all-atomic molecular dynamic studies of human CDK8 and provided insights into two point mutations D173A and D189N within the activation loop by the hydrogen bond (H-bond) dynamic study of the activation loop residues and the MMPB/SA method [28].

We have employed all atom unbiased molecular dynamics approach to understand the importance of CycC and the effect of the motions of the major CDK8 regions such as the activation loop and the αC helix on ligand binding. We studied 5 DMG-in CDK8 complexes and 5 DMG-out CDK8 complexes, and built homology models for the missing activation loop. We present here the new details of protein structural dynamics and ligand binding modes, confirm the importance of CycC to the stability of the ATP binding site and the allosteric site, and describe new findings that would be utilized to improve the existing ligands.

## 2. Methods

### 2.1 Structures of CDK8/CycC and the ligands

We performed MD simulations to study the dynamics of DMG-in and DMG-out apo CDK8/CycC proteins, five DMG-in and five DMG-out (structure-kinetic relationship series, SKR) CDK8/CycC-ligand complexes. The MD simulation indices were assigned to the studied systems for simplicity. They together with PDB IDs of the crystal structures used as initial conformations are listed in Table 1. We manually mutated the crystal structure 4F7N to yield the complex of CDK8/CycC-SKR10 whose crystal structure is not available. For the apo CDK8/CycC in DMG-in conformation, we studied two systems, one in the free state (4G6L) and the other in the bound state (5CEI) after the removal of the ligand 50R. For the apo CDK8/CycC in DMG-out conformation, we studied the one in the bound state (4F6W) after the removal of the ligand SKR1. The molecular structures of the 10 ligands are listed in Figure 1. We kept residues 1 to 359 for CDK8 and residues −2 to 257 for CycC for the MD simulations. To build the missing activation loop, we used p38 (PDB ID: 1W82) as the reference structure and constructed homology models of the CDK8 activation loop using SWISS-MODEL [29–31]. Then we aligned residues Asp173 and Arg200, and manually added the homology model of the activation loop to CDK8. We added the other missing loops by using SWISS-MODEL with the corresponding crystal structures as the references.

**Figure 1.**
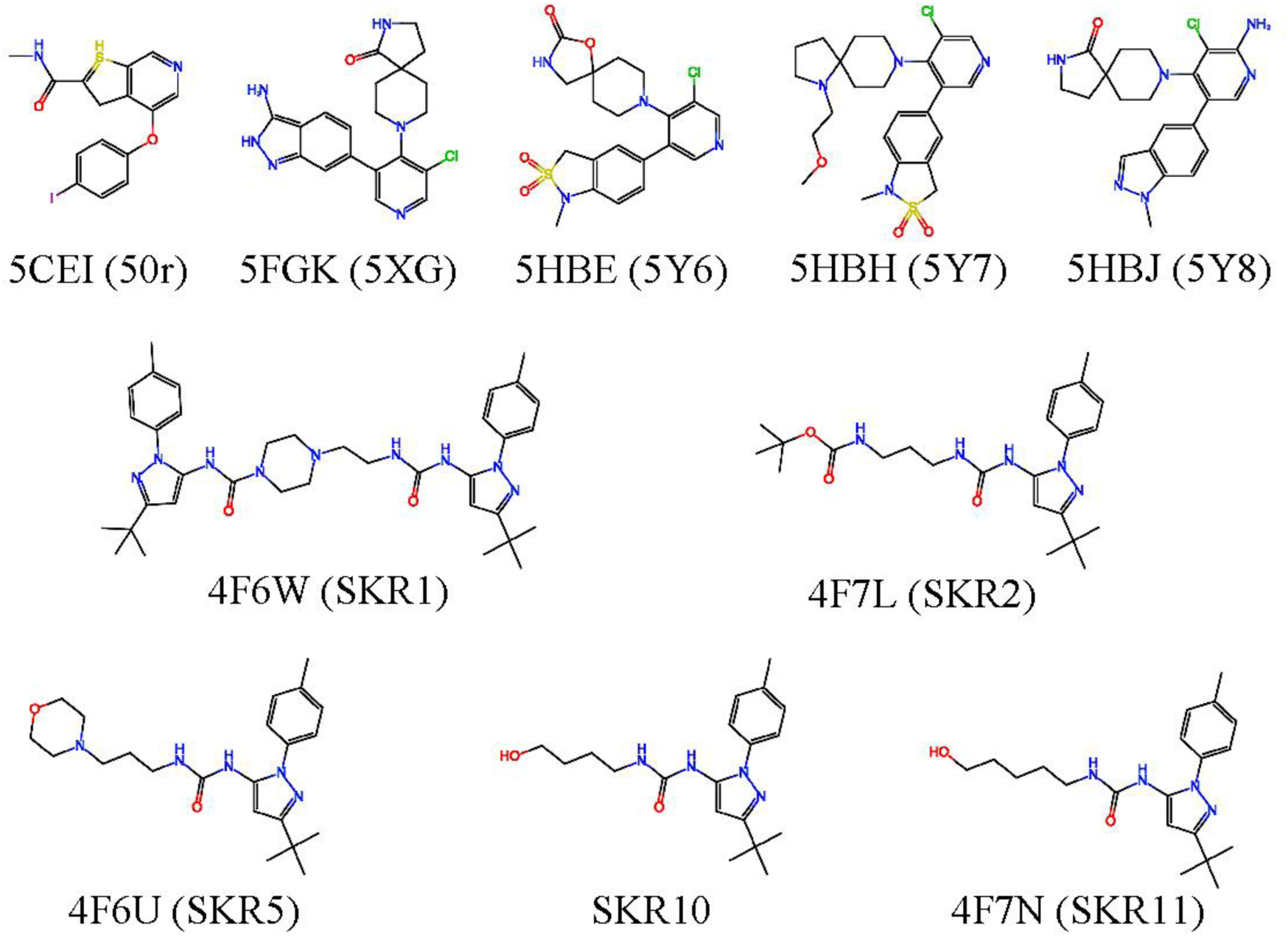
The molecular structures of the 10 compounds used in this work.

**Table 1.**
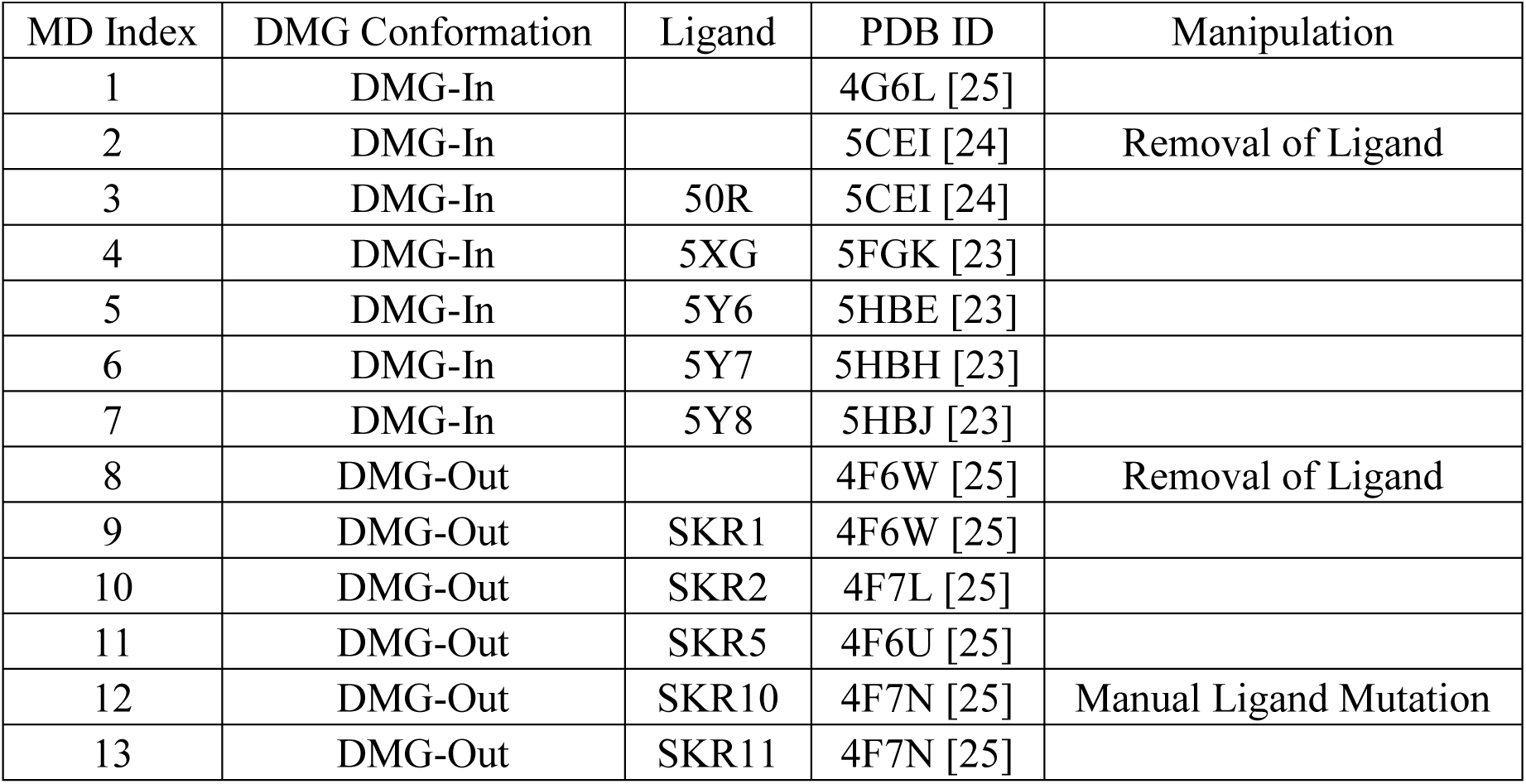
The list of 13 CDK8/CycC systems studied in this work. The references to the crystal structures are provided with PDB IDs.

### 2.2 Unbiased Molecular Dynamics (MD) simulation

The Amber 14 package with an efficient GPU implementation [32–34] was employed for the MD simulations of dynamic processes in the binding between CDK8/CycC and the ligands. Amber 99SB and General Amber Force Field (GAFF) [35–37] were applied to CDK8/CycC, and the ten ligands, respectively. Single protonation states were used for all histidine residues according to protonation state predicted for CDK8/CycC complex using MCCE [38, 39]. 6 Cl-ions were placed to maintain a neutral system. Minimization on the hydrogen atoms, side chains and the entire protein complex was performed for 500, 5000 and 5000 steps, respectively. After being solvated with a rectangular TIP3P water box [40], the edge of the box is at least 12 Å away from the solutes. The system went through a 1000-step water and 5000-step system minimization to correct any inconsistencies. Then we equilibrated the water molecules with the solutes fixed for 20 ns at 298K in an isothermic-isobaric (NPT) ensemble. Next, we relaxed the system by slowly heating it during an equilibrium course of 10 ps at 200, 250 and 298 K. We performed production run in an NPT ensemble with a 2-fs time step. The Langevin thermostat [41, 42], with a damping constant of 2 ps^-1^, was used to maintain a temperature of 298 K. The long-range electrostatic interactions were computed by the particle mesh Ewald method [43] beyond 8 Å distance. The SHAKE algorithm [44] was used to constrain water hydrogen atoms during the MD simulations. We performed 500 ns of MD production runs on each complex and apo protein by using CPU parallel processing and local GPU machines. We collected the resulting trajectories every 2 ps and resaved the trajectories for analysis at intervals of 20 ps.

### 2.3 Analysis methods

#### (1) RMSD

Root-mean-square deviation (RMSD) of a group of atoms in a molecule with respect to a reference structure is calculated as,

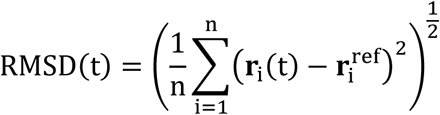

where *n* is the number of atoms in the group, ***r**_i_(t)* and 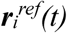 are the Cartesian coordinates of the atom *i* at time *t* in the trajectories and in the reference structure respectively. We used the conformation in the corresponding crystal structure after adding the missing loops as the reference and then aligned the trajectories to the reference. We only considered heavy atoms in the trajectories alignment. RMSD was computed using VMD [45].

#### (2) RMSF

The root-mean-square fluctuation (RMSF) is a measure of the deviation between the position of a particle and its average position. We computed RMSF for the α-carbon atom of each residue in CDK8 as:

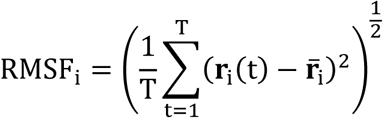

where *T* is the total MD time, ***r**_i_(t)* and ***r**_i_* are the Cartesian coordinates of the α-carbon atom in residue *i* at time *t* and the its average over time *T*. We used the same aligned trajectories to compute RMSF as in the RMSD calculation.

#### (3) Cartesian PCA

We performed classical principal component analysis (PCA) [46–48] on α-carbon atoms in the 500 ns trajectories saved every 20 ps (25000 frames in total). To obtain PC motions in different periods of MD simulations, we divided the aligned 500 ns trajectories of each system into five 100 ns short ones and performed PCA on α-carbon atoms in Cartesian coordinates with each of the five short trajectories. The average positions of the α-carbon atoms were used as reference to compute the covariance matrix. The first PC modes were saved and analyzed.

#### (4) Hydrogen Bonding analysis

In this work a H-bond (X-H…Y) is considered formed if the distance between H and Y is smaller than 2.5 Å and the complimentary angle of X-H…Y is smaller than 30° (Figure SI-1). We used an in-house script to process through the trajectories for direct H-bonds between ligands and CDK8 as well as mediating water molecules that connect ligands and CDK8. H-bonds between ligands and the same residues are merged into one residue-ligand H-bond formation. The occurrence percentage of a H-bond is calculated as the number of the frames where the H-bond is found divided by the total frames (25000).

#### (5) Residue-wise interaction

We computed the ligands interactions with the 619 CDK8/CycC residues. For each residue we computed the sums of van der Waals, Coulombic, Generalized Born (GB) energy terms for the ligand with the residue (E_L+R_), the ligand alone (E_L_), and the residue alone (E_R_), and then computed the interaction energy Δ E = E_L+R_ - E_L_ - E_R_. Only residues that directly interact with the ligands are considered in our analysis.

#### (6) MMPB/SA

We used the MMPB/SA method [49] to evaluate the ligand binding affinities to CDK8/CycC. MMPB/SA estimates the energy (E) of a system which can be either protein(P), ligand(L) or complex(PL):

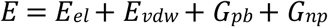

where *E_el_* and *E_vdw_* are electrostatic and van der Waals energy, G_pb_ is the solvation energy computed by solving the Poisson Boltzmann (PB) equation, G_np_ is the nonpolar energy estimated from solvent accessible surface area. The binding energy ΔE is estimated by averaging the energy difference computed by subtracting the energy of the free protein (E_P_) and the free ligand (E_L_) from the energy of the complex (E_PL_) as,

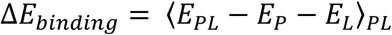

#### (7) Volume analysis

We evaluated the volume of the ATP binding site. We applied a grid box with a spacing of 1 Å to each conformation of a MD trajectory. The position of this grid box was decided by the concurrent minima and maxima of the Cartesian coordinates of the α-carbons of CDK8 residues 66, 30, 171, 27 and 156 of each conformation. Then we computed the solvent accessible volume in the grid box as the volume of the ATP binding site.

#### (8) T-Analyst

We performed T-Analyst [50] on the CDK8 protein using the CDK8/CycC-Ligand complex trajectories. Only psi and phi angles were considered in the correlation matrix. We also computed the correlation maps with the residues grouped according to the secondary structure of CDK8, where separate α-helices, β-strands and loops were considered as individual groups.

## 3. Results and Discussion

### 3.1 Binding Pocket and Ligand Binding Modes

Our MD results about the binding modes, the major residues that form H-bonds with ligands, and the major residues that contribute the ligand-binding interactions are presented in Figure 2, Table 2 and Table 3, respectively. Type I ligands occupy the ATP binding pocket and form consistent interactions with ALA100, LYS52 and ARG356. Figure SI-2 presents the binding mode of 50R as an example. Ligand 50R forms a stable H-bond with ALA100 through its nitrogen on the benzothiophene ring. The occurrence percentage of this H-bond is 64%. Its amide moiety interacts with LYS52 by a H-bond which has an occurrence percentage of 38%, and its benzene ring forms a cation-*π* interaction with ARG356. The major residues that contribute van der Waals (vdW) interactions with 50R are LEU158, ARG356 and VAL35. This binding mode of 50R is in good agreement with the crystal structure 5CEI and is retained through the 500 ns MD run. Ligand 5XG finds a cation–π interaction of its indazole phenyl ring with ARG365, binds to the hinge by its pyridine ring, and forms a H-bond with LYS52 by the 2,8-diazaspiro[4.5]decan-1-one moiety. In addition, its 3-aminoindazole moiety forms a highly stable H-bond with VAL27 with an occurrence percentage of 78%. The major vdW contributors are LEU158, ARG356, VAL35 and TYR32. This binding mode of 5XG keeps consistent with the crystal structure and stable in the course of 500 ns MD run. Ligand 5Y6 has a similar binding mode as 5XG, except that it loses the interaction with VAL27 but picks up contacts with TYR32 and ASP173. The occurrence percentages of the H-bonds with TYR32 and ASP173 are both below 10%, and they are not found in the crystal structure. The major vdW contributors are LEU158, ARG356, VAL35 and TYR32.

**Figure 2.**
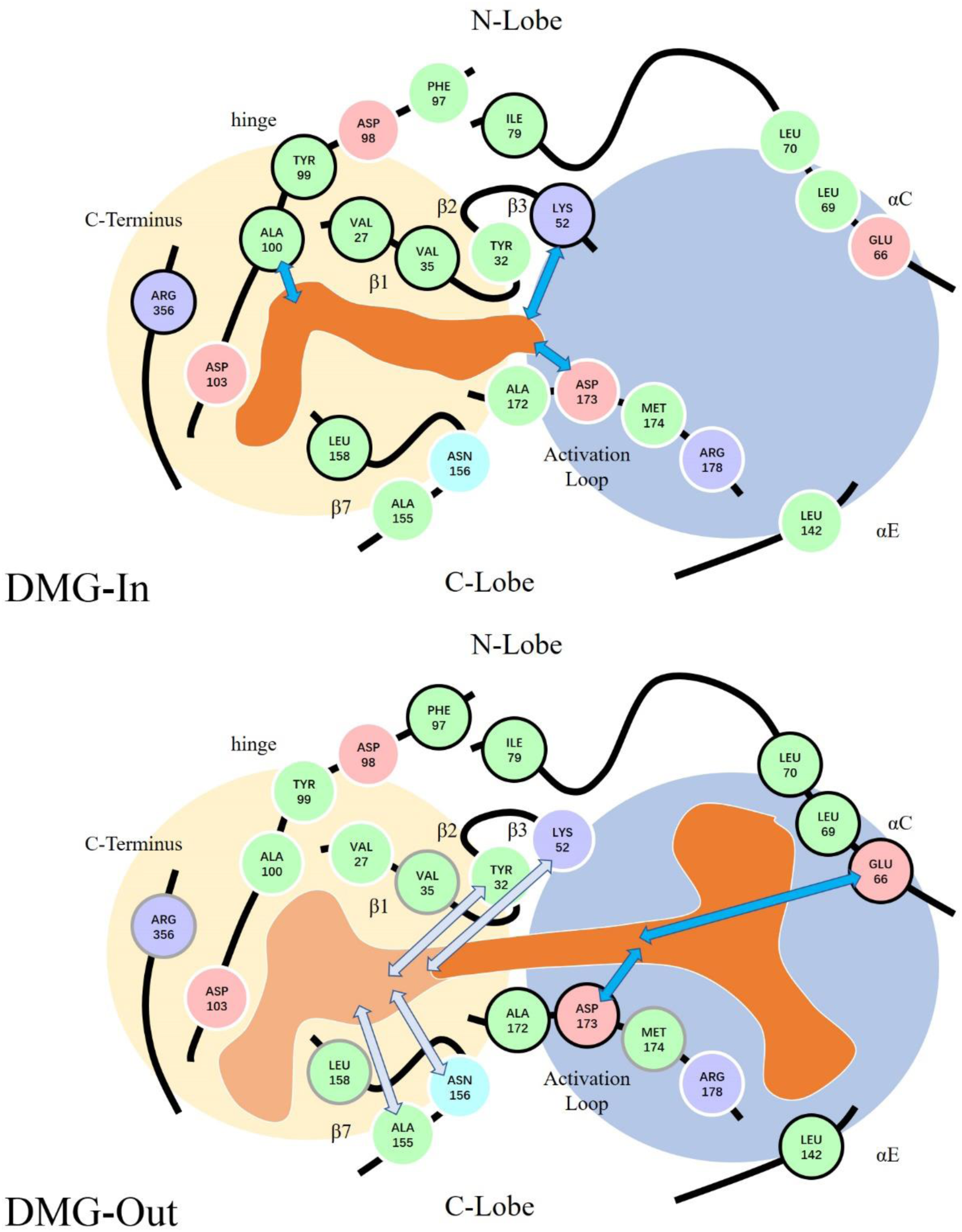
Illustration of ligand binding modes in DMG-in and DMG-out conformations. The regions of CDK8 are indicated by black lines, and labeled correspondingly. Important residues are shown by spheres, where non-polar residues are in green, polar and neutral residues are in cyan, positive charged residues are in blue and negatively charged residues are in red. Residues with interactions stronger than −1 kcal/mol with DMG-in/DMG-out ligands are blacked bordered. Residues with interaction stronger than −1 kcal/mol with SKR1 are gray bordered. Other residues are white bordered. Important hydrogen bonds between residues and ligands are indicated by deep cyan arrows, while hydrogen bonds with big DMG-out ligands are indicated by light cyan arrows. Ligand binding modes are shown by orange areas, and light orange area indicates additional binding area by SKR1. The two big circles indicate the ATP binding site (light orange) and the allosteric binding site (light blue).

**Table 2.** The major residues that form hydrogen bonds with the type I and type II ligands and their occurrence percentages. Hydrogen bonds with percentage smaller than 1% in 500 ns are not shown. Universal hydrogen bonding residues are highlighted.

| Type I |  |  |  |  |  |
| --- | --- | --- | --- | --- | --- |
| MD Index | 3 | 4 | 5 | 6 | 7 |
| Ligand | 50R | 5XG | 5Y6 | 5Y7 | 5Y8 |
| PDB ID | 5CEI | 5FGK | 5HBE | 5HBH | 5HBJ |
| VAL27 |  | 79.28% |  |  |  |
| TYR32 |  | 25.76% |  | 4.12% | 5.94% |
| LYS52 | 40.00% | 48.00% | 40.00% | 42.48% | 38.80% |
| GLU66 | 13.60% | 12.94% | 13.60% | 2.92% | 2.98% |
| ASP98 |  |  |  |  | 51.77% |
| ALA100 | 64.03% | 56.76% | 64.03% | 76.53% | 76.86% |
| ASP103 |  | 44.17% |  | 32.57% | 23.26% |
| HIE106 |  | 2.03% |  | 65.61% | 2.16% |
| ALA155 |  | 23.42% |  |  |  |
| ASP173 | 24.78% | 36.73% | 24.78% | 21.83% | 29.80% |
| ARG356 |  |  |  | 50.26% |  |
| Type II |  |  |  |  |  |
| MD Index | 9 | 10 | 11 | 12 | 13 |
| Ligand | SKR1 | SKR2 | SKR5 | SKR10 | SKR11 |
| PDB ID | 4F6W | 4F7L | 4F6U | 4F7N | 4F7N |
| VAL27 | 11.57% |  |  |  |  |
| TYR32 | 30.63% |  |  |  | 1.46% |
| LYS52 | 40.36% | 2.42% |  | 2.34% | 2.00% |
| GLU66 | 96.04% | 93.92% | 91.82% | 94.26% | 89.26% |
| ASP98 |  | 24.84% | 1.80% | 39.71% | 35.58% |
| ALA100 |  | 12.38% | 16.66% | 2.29% | 15.86% |
| ALA155 | 38.96% |  |  |  |  |
| ASN156 | 30.51% |  |  |  |  |
| ASP173 | 82.92% | 92.97% | 76.24% | 88.95% | 87.04% |
| ARG178 |  |  | 17.04% |  |  |
| ARG356 | 10.45% |  |  |  |  |

**Table 3.** The major residues that have interaction energies stronger than −1.0 kcal/mol with type I and type II ligands. All values are in kcal/mol.

| Type I |  |  |  |  |  |
| --- | --- | --- | --- | --- | --- |
| MD Index | 3 | 4 | 5 | 6 | 7 |
| Ligand | 50R | 5XG | 5Y6 | 5Y7 | 5Y8 |
| PDB ID | 5CEI | 5FGK | 5HBE | 5HBH | 5HBJ |
| LEU158 | -2.7±0.6 | -2.8±0.6 | -2.7±0.6 | -1.2±0.6 | -2.7±0.6 |
| ARG356 | -2.5±0.6 | -2.7±0.7 | -2.7±0.7 | +3.6±1.5 | -3.2±0.6 |
| VAL35 | -2.4±0.5 | -2.4±0.6 | -2.5±0.7 | -1.9±0.5 | -2.2±0.6 |
| LYS52 | -2.0±1.2 | -3.0±1.1 | -2.7±0.9 | -1.2±1.0 | -2.6±1.2 |
| TYR99 | -1.7±0.4 | -1.8±0.4 | -1.6±0.4 | -1.9±0.5 | -2.0±0.5 |
| ALA100 | -1.2±0.7 | -1.2±0.7 | -1.4±0.6 | -1.0±0.7 | -1.5±0.5 |
| PHE97 | -1.0±0.4 | -1.0±0.4 | -1.1±0.3 | -0.7±0.3 | -0.7±0.3 |
| TYR32 | -0.6±0.6 | -2.3±1.0 | -2.8±1.0 | -2.2±0.9 | -2.0±1.0 |
| VAL27 | -1.0±0.5 | -1.7±0.7 | -0.9±0.5 | -1.0±0.6 | -0.9±0.5 |
| ILE79 | -0.8±0.4 | -1.1±0.4 | -1.1±0.4 | -1.1±0.4 | -0.8±0.5 |
| ASP103 | +0.1±0.6 | +1.2±0.7 | +1.9±0.8 | -1.3±1.7 | +1.4±0.6 |
| Type II |  |  |  |  |  |
| MD Index | 9 | 10 | 11 | 12 | 13 |
| Ligand | SKR1 | SKR2 | SKR5 | SKR10 | SKR11 |
| PDB ID | 4F6W | 4F7L | 4F6U | 4F7N | 4F7N |
| GLU66 | -5.2±1.1 | -5.4±1.2 | -4.6±1.4 | -5.2±1.1 | -4.4±1.4 |
| MET174 | -3.3±0.8 | -2.8±1.0 | -1.1±0.8 | -1.4±0.8 | -0.7±0.8 |
| ARG356 | -2.8±0.9 | -0.1±0.5 | 0.0±0.4 | 0.0±0.3 | 0.0±0.3 |
| LEU70 | -2.7±0.8 | -2.7±0.8 | -2.8±0.8 | -2.7±0.7 | -2.9±0.8 |
| LEU158 | -2.4±0.8 | +0.9±0.6 | -0.6±0.6 | 0.0±0.4 | 0.0±0.6 |
| ASP173 | -2.0±1.0 | -2.3±1.0 | -1.8±1.0 | -2.2±1.0 | -1.7±1.0 |
| PHE97 | -1.9±0.7 | -2.2±0.8 | -2.3±0.7 | -1.4±0.8 | -1.7±0.8 |
| VAL35 | -1.9±0.7 | -1.7±0.8 | -1.0±0.6 | -0.1±0.6 | -0.3±0.7 |
| ALA172 | -1.7±0.8 | -1.7±0.9 | -2.0±0.7 | -1.7±0.7 | -1.6±0.7 |
| ILE79 | -1.7±0.8 | -2.1±0.9 | -2.2±0.8 | -1.5±0.8 | -1.8±0.9 |
| LEU69 | -1.1±0.7 | -1.0±0.7 | -1.0±0.6 | -1.0±0.7 | -1.0±0.6 |
| ARG178 | 0.0±0.1 | 0.0±0.3 | -1.2±1.0 | 0.0±0.2 | 0.0±0.4 |
| LEU142 | -0.9±0.5 | -1.0±0.6 | -0.9±0.6 | -1.0±0.6 | -0.9±0.7 |

Ligand 5Y7 adopts a different binding mode than the crystal structure from roughly 40 ns. In this conformation, the salt bridge between LYS52 and ASP173 disappears, resulting in the breathing motion of the CDK8 N-lobe and C-lobe and the opening of the binding site. Interestingly, although 5Y7 still has a cation–π interaction with ARG356, but the energy contribution from ARG356 becomes positive, i.e., repulsive interaction. It is because the opening of binding site makes ARG356 to be exposed to the solvent, and the solvation energy term contributes to the repulsive interaction between ARG356 and 5Y7. 5Y7 forms a H-bond with ALA100 via nitrogen in the 5-membered ring with an occurrence percentage of 76%. It also experiences H-bonding with HIS106 via its SO2 group with an occurrence percentage of 65%. Recurring flipping of the pyrrolidine ring in 5Y7 at times enables a H-bond with LYS52 via the oxygen in its methoxyethyl moiety, with an occurrence percentage of 42%. The major residues that provide vdW contributions are LEU158 and TYR32. The pyrrolidine ring appears to be very flexible. The opened binding site allows the nitrogen on this ring to inverse several times in the 500ns MD run, and causes its attached pendant methoxyethyl moiety to swing near the αC helix (Figure 3). Interestingly the N-inversion correlates with the motion of the activation loop to some extent (Figure 4).

**Figure 3.**
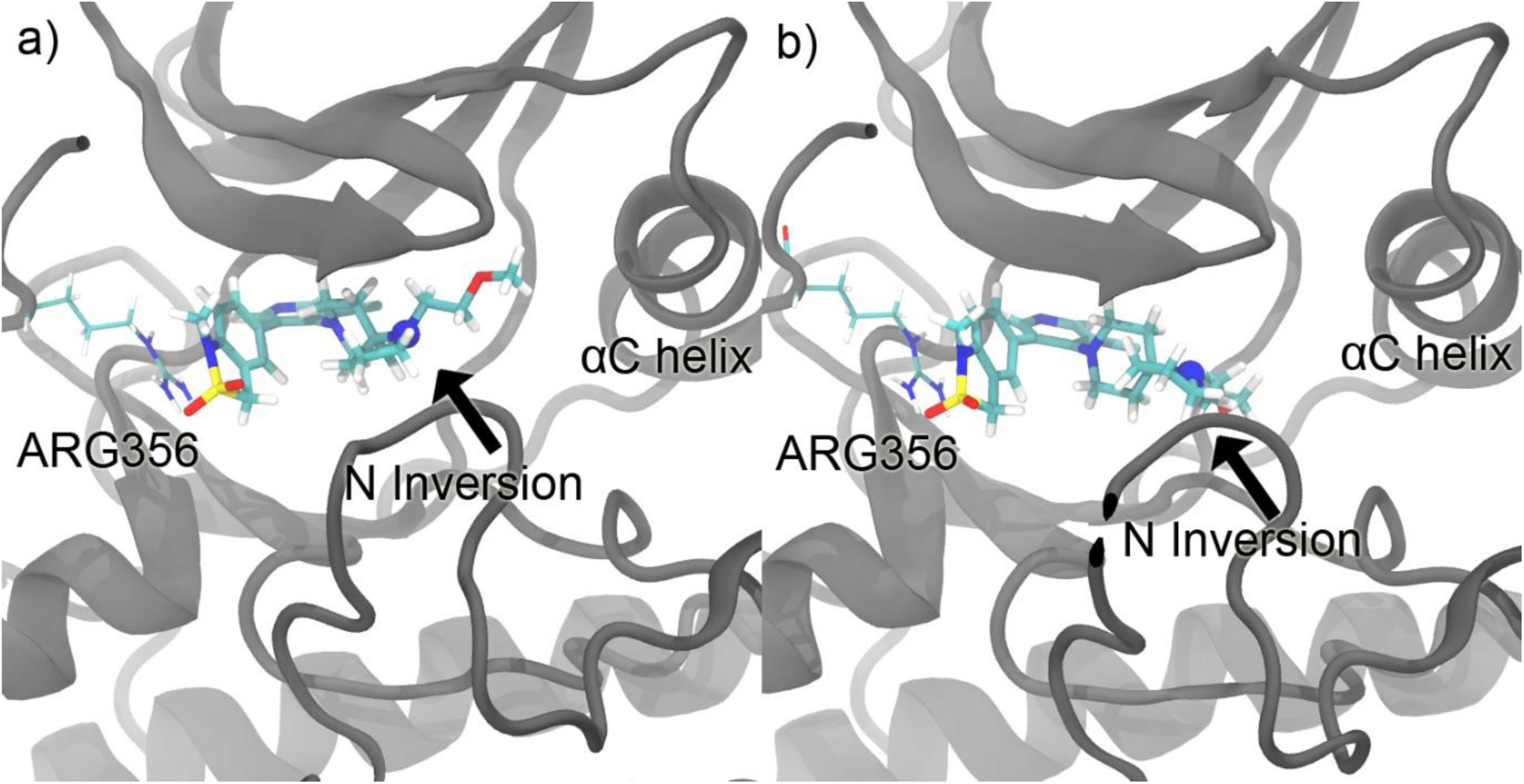
Flipping of the the pyrrolidine ring on 5Y7 in the course of 500 ns MD run.

**Figure 4.**
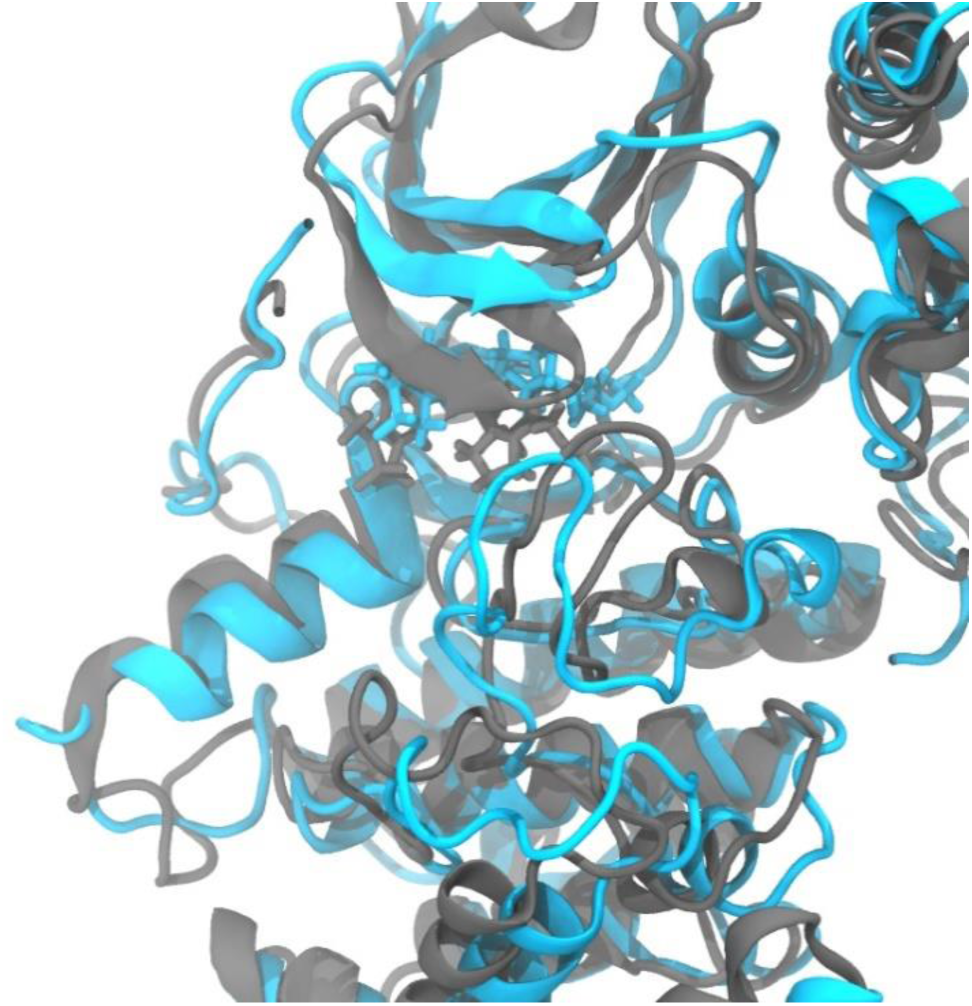
Superposition of two frames at 0 ns and 500 ns for the complex of CDK8/CycC and 5Y7. Gray: 0 ns. Cyan: 500 ns.

Ligand 5Y8 also forms its most stable H-bond with ALA100 like other DMG-in ligands via the nitrogen on the pyridine ring, with an occurrence percentage of 77%. Its second-most stable H-bond is formed between ASP98 and the amine group on the pyridine ring, with an occurrence of 52%. With small conformational changes in the protein, LYS52 moves close enough to 5Y8 to experience H-bonding, with the ligand’s carbonyl group acting as the acceptor. This bond has an occurrence percentage of 31%. Similar to 5Y6, the benzene ring on 5Y8 interacts with ARG356 via cation-π interaction. The major residues to provide vdW contributions are LEU158, ARG356, and TYR32.

The binding of type II ligands to CDK8 features the very stable H-bonding network involving the urea linker on the ligands, ASP173 and GLU66 at the allosteric site. This network is further stabilized by the strong salt bridge between GLU66 and LYS52. Type II ligands also extend to the ATP binding site to gain additional strength. In general, the binding modes predicted by our MD simulations highly resemble those in the crystal structures [refer to Figure 3 in 25]. SKR1 has the largest size among the 5 type II ligands. Since both the allosteric site and the ATP binding site are hydrophobic, SKR1 has extensive vdW interactions with its surrounding residues. Its benzene ring at the ATP binding site interacts with ARG356 through cation-π stacking. Its H-bonds with ASP173 and GLU66 are highly stable, with the occurrence percentages of 83% and 96% respectively. But it doesn’t have contacts with the hinge region. SKR2 has the similar binding mode as SKR1. Since it has a smaller structure than SKR1 in the part that occupies the ATP binding site, it has less vdW contacts in this region as compared with SKR1. Instead of cation-π, it has vdW interaction with ARG356. For SKR5, besides the key H-bonding network with ASP173 and GLU66, its terminal [3-(morpholine-4-yl)propyl] group forms another H-bond with ALA100 in the hinge region. Although the occurrence percentage of this H-bond is as low as 17%, it does provide SKR5 detectable residence time [25]. Due to the short length of its carbon chain, SKR10 stretches to the ATP binding site but could only make an occasional contact with ASP98 through a H-bond, the occurrence percentage of which is poorly 2%. In contrast, SKR11 with one more carbon on its carbon chain than SKR10, is able to form a slightly more stable H-bond with ASP98, with an occurrence percentage of 10%. SKR11 also interacts with ALA100 occasionally through an unstable H-bond.

We calculated the binding energies of the ligands using MMPB/SA method. The computed values are grouped by DMG-in and DMG-out conformations and their correlations with experimental values are plotted in Figure 5. Type I ligands have much better MMPB/SA-Experiment relationship than type II ligands. The MMPB/SA energies in our calculations don’t include the entropy term. The structures of type I ligands are more rigid than those of type II ones and thus the influence of the entropy term on the binding energies of type I is smaller than that of type II. Therefore, MMPB/SA provides more accurate predictions on type I ligands. The energy breakdowns can be found in the supplement materials (Table SI-1). The binding is mainly driven by vdW interactions for both type I and type II ligands. 5XG and 5Y6 are the top 2 potent ones among the 5 type I ligands. Since all the type I ligands have roughly similar overall electrostatic contribution to the binding, it is the vdW interactions that make the difference in potency. The residue-wise interaction analysis indicates that both 5XG and 5Y6 receive strong vdW contributions from TYR32, VAL35, LEU158 and ARG356. In contrast, VAL35 and ARG356 don’t contribute to 5Y7, leading to much less vdW contribution to this ligand. Moreover, ARG356 applies an energy penalty on the binding of 5Y7, indicating that this ligand doesn’t fit the binding site very well. These results suggest that complementary fit may be the key to improve the potency of CDK8 type I ligands.

**Figure 5.**
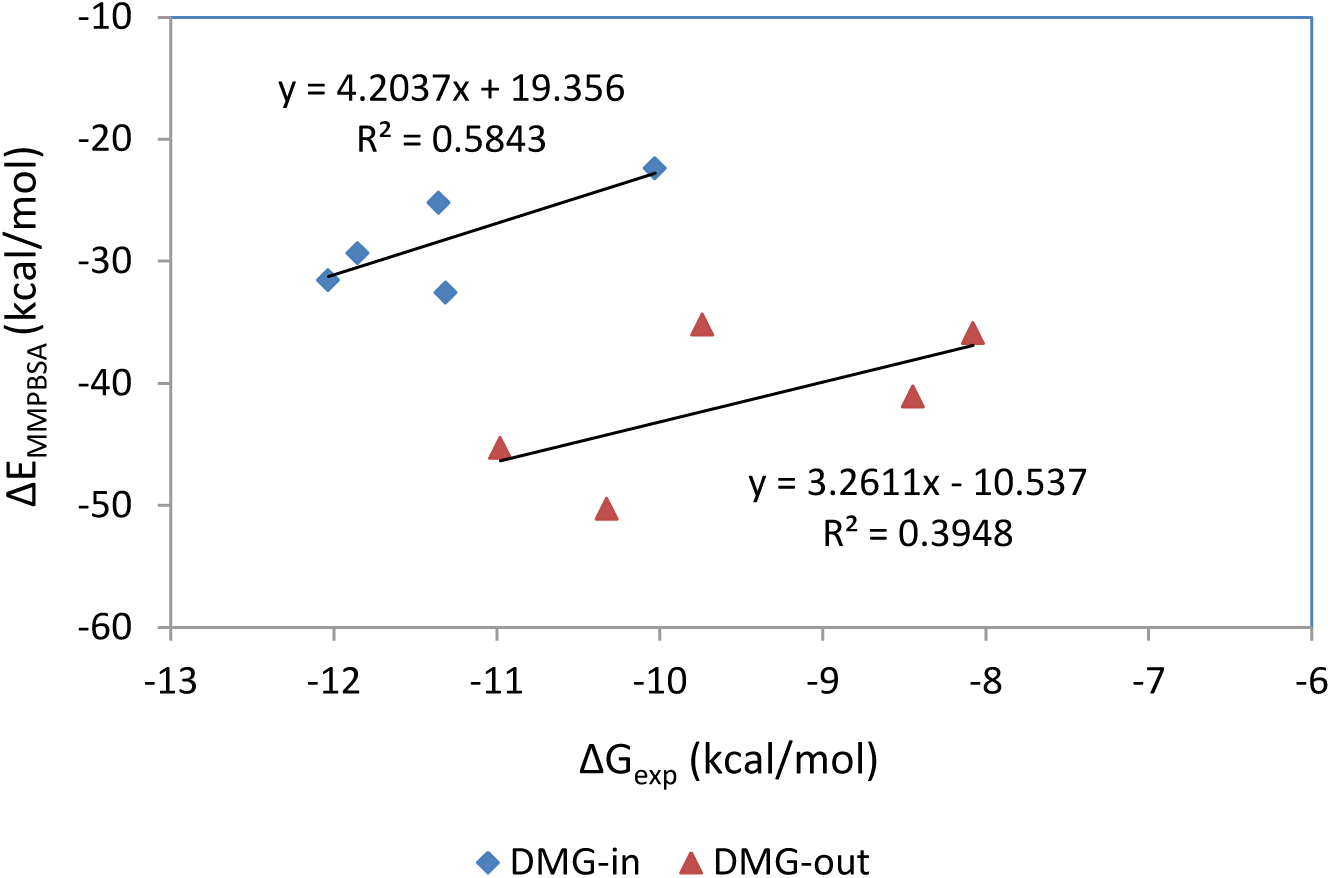
The relationship between the experimental free energies and the calculated MMPB/SA energies.

The contributing factors of type II ligand binding are more complicated than type I ligands. Type II ligands share the same scaffold that binds to the allosteric binding site of CDK8, but have different structures that extend into the ATP binding site. SKR10 and SKR11 have less bulky structures than SKR1, SKR2 and SKR5, and thus have less favorable vdW interactions with the ATP binding site. Electrostatic interactions play a more important role in the binding of SKR10 and SKR11. Moreover, the MMPB/SA suggests that SKR1 has stronger binding energy than SKR2, but experimental data favors SKR2 by 0.5 kcal/mol. Since entropy contribution was not considered in our MMPB/SA calculations, entropy could account for this discrepancy. As compared with SKR1 the smaller SKR2 is less confined and retains more freedom, and therefore SKR2 suffers from less entropic penalty. In addition, RMSF result shows that CDK8 bound with SKR2 is more flexible than when bound with SKR1, suggesting that protein also pays less entropy penalty with SKR2 (Figure 6).

**Figure 6.**
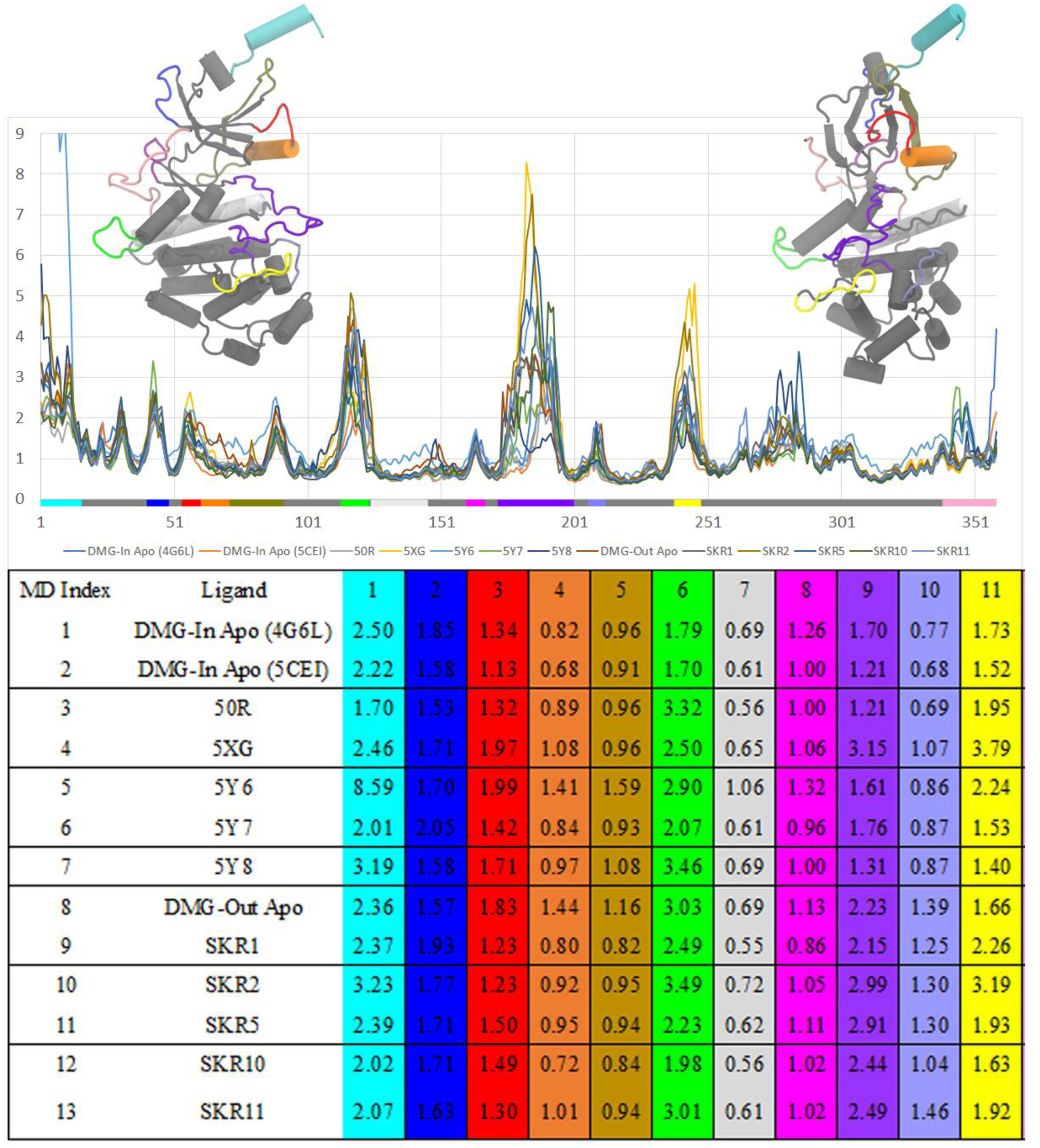
RMSF for all the studied CDK8/CycC systems. The color bar on the x-axis marks the secondary structure of CDK8. The table beneath the plot lists the 12 regions with large motions and highlighted with colors that correspond to the color bar.

### 3.2 Major CDK8 Motions

RMSF analysis on the MD frames reveals the regions that have major motions. Figure 6 presents the RMSF for all the 13 CDK8/CycC systems studied in this research. This figure shows that all the large motions occurred roughly at the same regions in all 13 systems. The table in this figure summarizes the regions with large motions (peaks) as presented in the plot. Different ligands cause fluctuations to different extents in some major regions. For the activation loop (region 9 in the table), the RMSF is 1.21 with 50R and 3.15 with 5XG. However, such large discrepancy in RMSF of the activation loop doesn’t affect the binding modes of these two ligands. They keep the poses resembling the crystal structures in the course of 500 ns MD runs, presumably because the large motion of the activation loop comes from its part exposed to solvent and had little influence on the binding pocket. In general, the activation loop is more flexible in the DMG-out conformation than in the DMG-in conformation. This is also reflected in the crystal structures, where in DMG-in conformations, the missing residues of the activation loop are less than in the DMG-out conformations. For the αC helix (region 4), type II ligands stabilize it because the urea linker on the type II ligands forms two H-bonds with GLU66, a residue that is absolutely conserved in the αC helix [27], and GLU66 forms a salt bridge with LYS52. This H-bonding network appears to be quite stable in the MD run – the occurrence percentage of the H-bonds between ligands and GLU66 is constantly higher than 90%. Thanks to the H-bond with LYS52, type I ligands have indirect connection with the αC helix. But this H-bonding network is much less stable than that of type II ligands – the occurrence percentage of the H-bond between ligands and LYS52 is below 50%. As a result, the αC helix is more flexible in the DMG-in conformation than in the DMG-out conformation.

It is worth noting that the αB helix (region 1) in MD5 has remarkably large motion, as shown in Figure 7. In the crystal structure 5HBE the N-terminal αB helix of CDK8 is found to have contacts with the CycC H5 helix and make a significant contribution to the binding between CDK8 and CycC [27]. The conformations in MD5 resemble the crystal structure 5HBE in the first 300ns MD run. The H-bonds between ARG13^CDK8^ and GLU144^CycC^ remain stable in this period. Afterwards the αB helix swings away. It still keeps contacts with the CycC H5 helix through the H-bonds between ARG13^CDK8^ and ASP147^CycC^ It also finds new contacts with the short H_N1_ helix through TYR3^CDK8^ and ASP20^CycC^ This new pose is further stabilized by the salt bridge between GLU12 and ARG40 and stays in the rest of MD run. Interestingly among all 13 systems this is the only one in the 500 ns MD run that we observed this new conformation. Due to this conformation change the αC helix and the activation loop in the complex with ligand 5Y6 also have markedly larger motions than those of other type I ligands.

**Figure 7.**
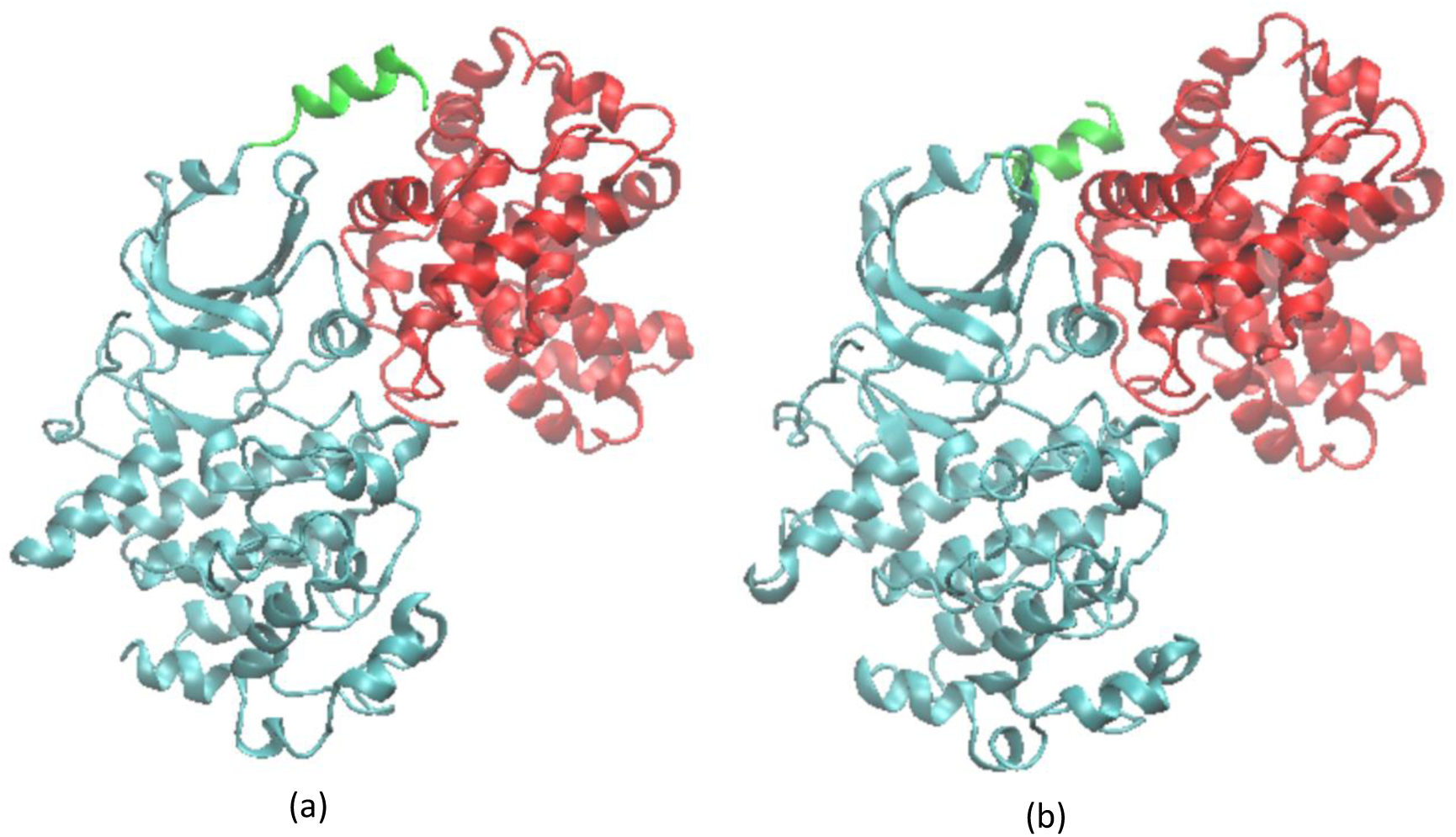
Conformational change of the αB helix in the 500 ns MD simulation. Cyan: CDK8; red: CycC; green: the αB helix. (a) Snapshot at 100 ns for the complex of CDK8/CycC and 5Y6. This MD structure is similar to the crystal structure 5HBE. (b) Snapshot at 360ns for the same complex. The αB helix adopts a different pose to bind with CycC.

The effect of ligand binding on the flexibility of CDK8 appears to be complicated. We studied two apo CDK8/CycC systems in the DMG-in conformation, one in the free state (MD1) and the other in the bound state with the removal of ligand 50R (MD2). The activation loop in MD2 has the exactly same RMSF value as its counterpart in MD3. The activation loops in the systems with other type I ligands have higher flexibility than MD2 but comparable with MD1. Therefore, it seems ligand binding has little influence on the flexibility of the activation loop in the DMG-in conformation. For the DMG-in conformation except region 12, the C-terminal tail, all the other regions with major motions have similar flexibility in both the free state and the bound state, as indicated by the RMSF values. ARG356 in region 12 forms cation-π interaction with type I ligands, resulting in much smaller motion in this region in the bound state. Therefore type I ligands stabilize the C-terminal tail. We studied one apo CDK8/CycC systems in the DMG-out conformation, the one in the bound state with the removal of ligand SKR1. The activation loop in DMG-out conformation is generally more flexible than in DMG-in conformation. However, its flexibility can become either higher or lower after ligand bind to the protein, indicating that ligand binding has no clear influence on it either. Type II ligands are found to stabilize the αC helix in the DMG-out conformation by the aforementioned mechanism.

The Cartesian principal component analysis (PCA) provides the information about protein thermal motions. Five major motions are observed in both the DMG-in and the DMG-out conformations, with/without ligands, as shown in Figure 8. (A) is breathing motion in which N lobe bends and unbends relative to C lobe about the hinge which connects the two. (C) is rotational motion in which the two lobes rotate back and forth relative to each other. (B) and (D) are the bending and rotational motions between CDK8 and CycC, respectively. (E) consists of the motions of the αB helix, the activation loop and the loop connecting αD and αE helixes. Motions of (A) through (D) have considerable impacts on protein-ligand binding and unbinding. A lot of attentions were paid to the side chain rearrangements which are indeed important to the ligand binding modes. However, the binding sites of proteins are usually enclosed areas, and the protein motions facilitate ligand binding by opening up the binding sites. The breathing motion was found to be closely related to the binding/unbinding pathways of p38 MAP kinase [51, 52]. Motion (E) may be related to the conversion between DMG-in and DMG-out conformations [53–55]. The effect of the motions on the ligand binding kinetics of CDK8 is worth further investigation.

**Figure 8.**
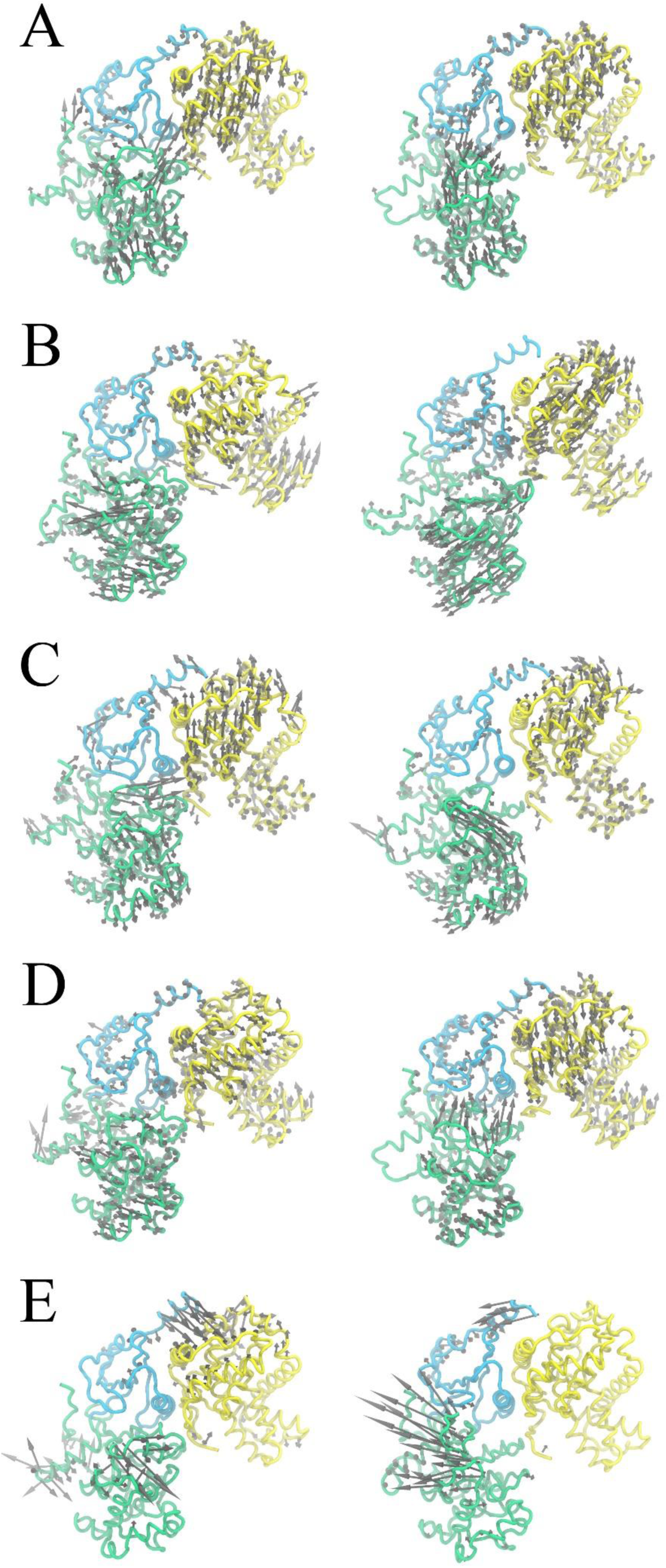
The first PC mode from Cartesian PCA of apo CDK8/CycC or CDK8/CycC-Ligand complexes using 100 ns trajectories. The breathing motion between N-lobe (cyan) and C-lobe (green) (A), breathing motion between CDK8 and Cyclin (yellow) (B), rotational motion between N-lobe and C-lobe (C), rotational motion between CDK8 and Cyclin (D), and loop motions (E) are indicated by gray arrows for DMG-In (Left) and DMG-Out (Right) conformations. The DMG-In PC modes use MD1 100-200 ns (A), MD3 0-100 ns (B), MD1 200-300 ns (C), MD5 0-100 ns (D), MD5 200-300 ns (E). The DMG-Out PC modes use MD12 100-200 ns (A), MD8 100-200 ns (B), MD12 300-400 ns (C), MD9 100-200 ns (D), MD10 100-200 ns (E).

In addition, different protein motions are observed in different periods in the 500ns MD simulations. The motion changes are resulted from the changes of the regions with the large RMSF values along the simulation time. For example, MD5 starts with (D), a rotational motion between CDK8 and CycC. Then the movement of the αB helix gradually picks up intensity. Between 300ns and 400ns, the protein motion mainly consists of the vibration of the αB helix, which corresponds to its conformation change. Once the conformation change is complete, the protein motion becomes (B), the breathing motion between CDK8 and CycC. MD12 starts with (C), with large vibration of the activation loop and the loop between αD and αE. Then the vibration of the loop between αD and αE dies down, so the protein motion changes to (A) between 100ns and 200ns. Afterwards the vibration of the loop between αD and αE becomes stronger again and so the protein motion changes back to (C). Such motion changes are not available from crystal structures which can only provide a snapshot of one particular conformational state. Therefore, to understand the dramatic changes in the binding sites associated with the dynamic behaviors of CDK8 upon binding and to depict a complete picture of protein-ligand interactions, MD simulations provide a more practical alternative than crystal structures.

The correlation among the major regional motions is further confirmed by the dihedral angle correlation analysis. Figure 9 shows correlation maps of the selected systems. These maps show that all the motions more or less correlate with one another. In general, motions in the DMG-out conformation, especially in the apo protein, have stronger correlations than those in the DMG-in conformation. The correlations are found among every region in CDK8 for the DMG-out conformation, while for the DMG-in conformation, the correlations are mainly found at the loop regions, especially the activation loop. Correlations in the regions of N-lobe of the DMG-in conformation are relatively weak. Besides the loops, the β sheets, i.e., β2 (30 to 37), β3 (49 to 59), β4-β5 (79 to 96), β7-β8 (154 to 172), are the major regions where internal correlations occurred. On the contrary, α-Helices are less correlated to the motions, especially α-E, α-F, αG, α-H, α-I, α-J, which are on the N-Lobe and on the back side of CDK8. Except for MD8, all other simulations show no correlations in the α-C helix. In addition, correlations in apo protein are stronger than those in complexes, indicating that ligand binding weakens the correlated motions among protein regions.

**Figure 9.**
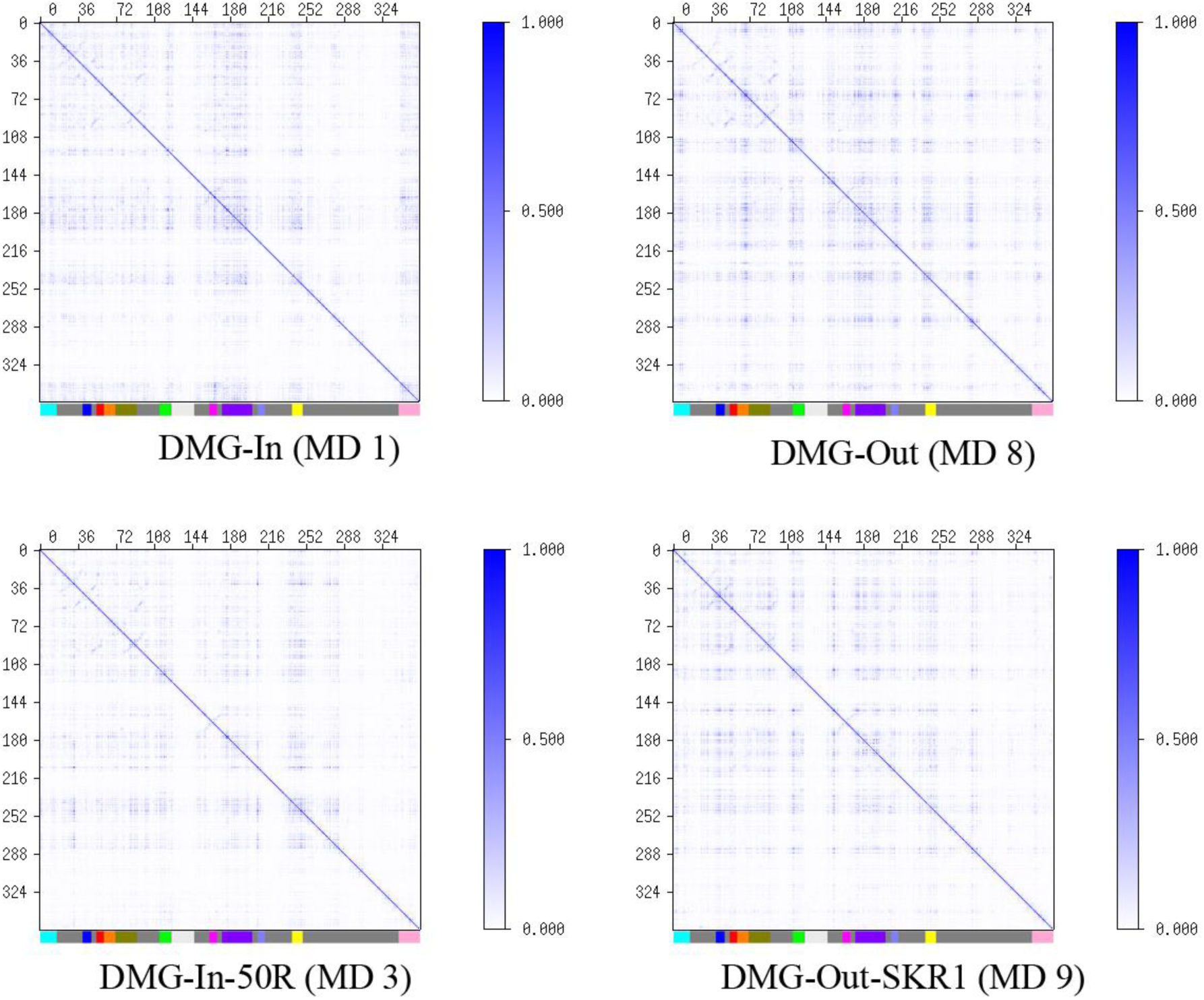
Correlation among the residue-wise motions obtained by dihedral angle analysis. Please refer to Figure 6 for the explanation of the color bar on x-axis.

### 3.2 Importance of Cyclin C on CDK8 and Ligand Binding

Cyclic C is critical to the functions of CDK8. To confirm the role of CycC in CDK8 and CDK8-ligand complex dynamics, we also performed MD simulations for CDK8-ligand complexes without CycC. Figure 10 shows the comparison of RMSF of CDK8 residues with and without CycC for the complex of CDK8 and 50R. Three regions have large discrepancies: the αB helix, the αC helix and the activation loop. In all of the three regions RMSF is significantly reduced in the presence of CycC. The αB helix is a part of the CycC binding interface. When CycC is bound to CDK8, the RMSD of the αB helix is reduced from around 20 Å to 4~6 Å, on the same level as other parts of the protein. But there is an exception. In the case of the complex of CDK8/CycC and 5Y6 (Figure 7), RMSD of the αB helix jumps from 5 Å to 25 Å after 300 ns. Å close inspection on the relevant frames revealed the binding mode change of the αB helix with CycC as aforementioned (refer to 3.2).

**Figure 10.**
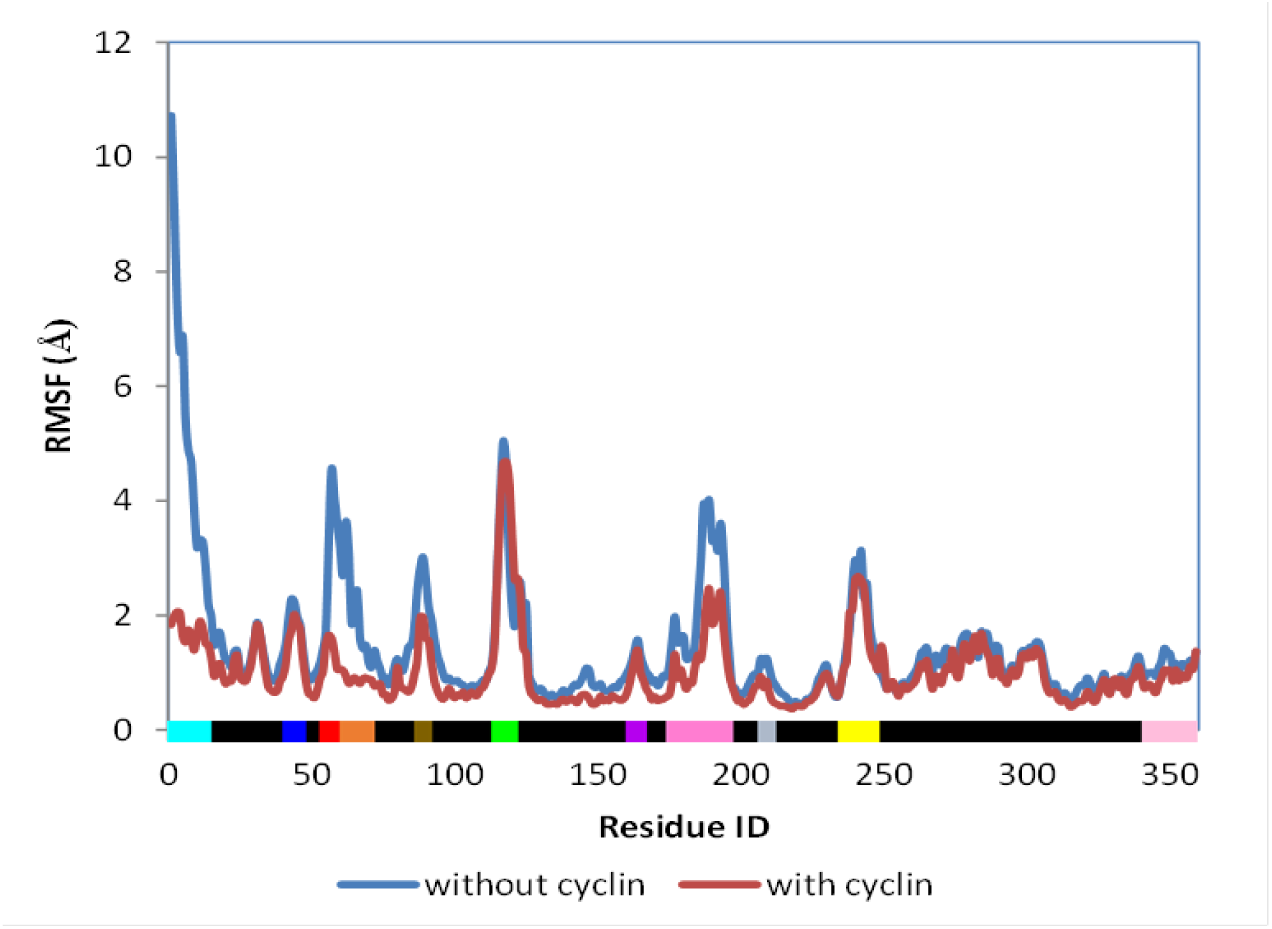
Comparison of RMSF of the complex of CDK8 and 50R with and without CycC.

The αC helix is adjacent to the binding site and has multiple conformations that could affect ligand binding. To determine the position of the αC helix, we measured the distance between the Cα carbon atoms of GLU66 and ASP173 [56]. Structures with a short DMG-αC-helix distance (4–7.2 Å) are marked αC-in, while structures with long distance (9.3–14 Å) are marked as αC-out. Structures with distances in between are marked as αC-out-like structures. Figure SI-3 presents an example. Without CycC this distance is usually found to fall into the αC-out conformation. The αC helix is found moving away from the binding site in both DMG-in and DMG-out conformations. In the case of apo protein the distance could get pretty large and greater than 14 Å. Figure 11 presents the conformation change of the αC helix in MD8 (apo CDK8 in DMG-out conformation) in the absence of CycC. The αC helix moves away from allosteric binding site and GLU66 moves out to solution. In the presence of CycC, however, αC-helix has much smaller fluctuation and the distance decreases to be in the αC-out-like category. Since GLU66, ASP173 as well as the αC helix are at the fringe of binding sites, shorter distance and smaller fluctuation helps the protein to make tighter and more stable contacts with ligands. Therefore, it is not surprising that ligands have larger motions and tend to have more binding modes when CycC is absent. Two major binding modes were observed in the complex of CDK8 and 5Y6, as shown in Figure 12. One resembles the crystal structure. In the other binding mode, the αC helix moves away from the ligand and the salt bridge between LYS52 and GLU66 is broken. This causes the β2-4 sheets above the binding site to move upward. These movements provide 5Y6 more room to explore the binding site. It still keeps a V-shape, but rotates by 90 degrees to pick up contacts with TYR32 and PHE97.

**Figure 11.**
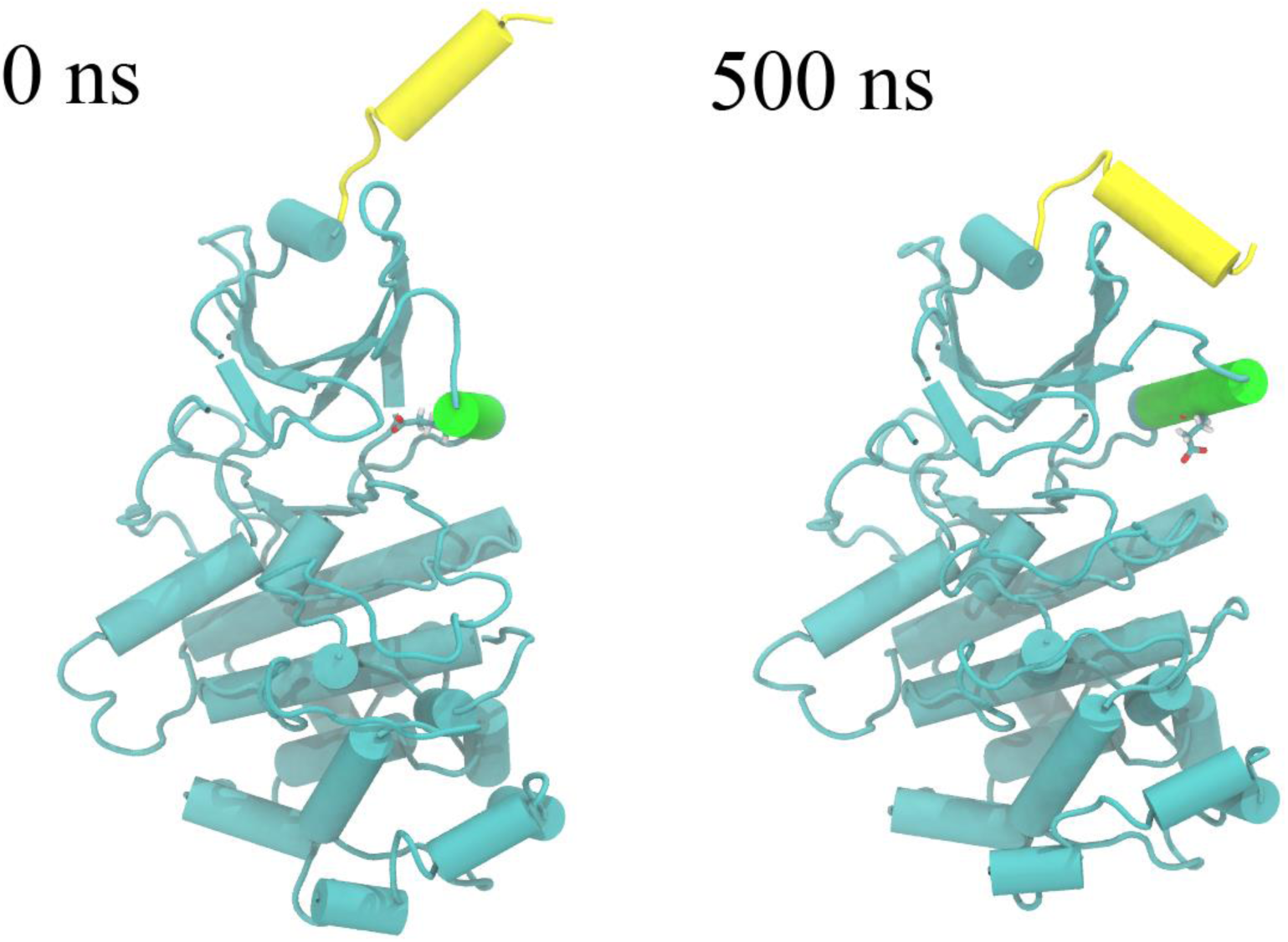
Conformation change of the αB and αC helixes in MD8 (apo CDK8 in DMG-out conformation) in the absence of CycC. The αB helix is in yellow, and the αC helix is in green. GLU 66 is shown in licorice.

**Figure 12.**
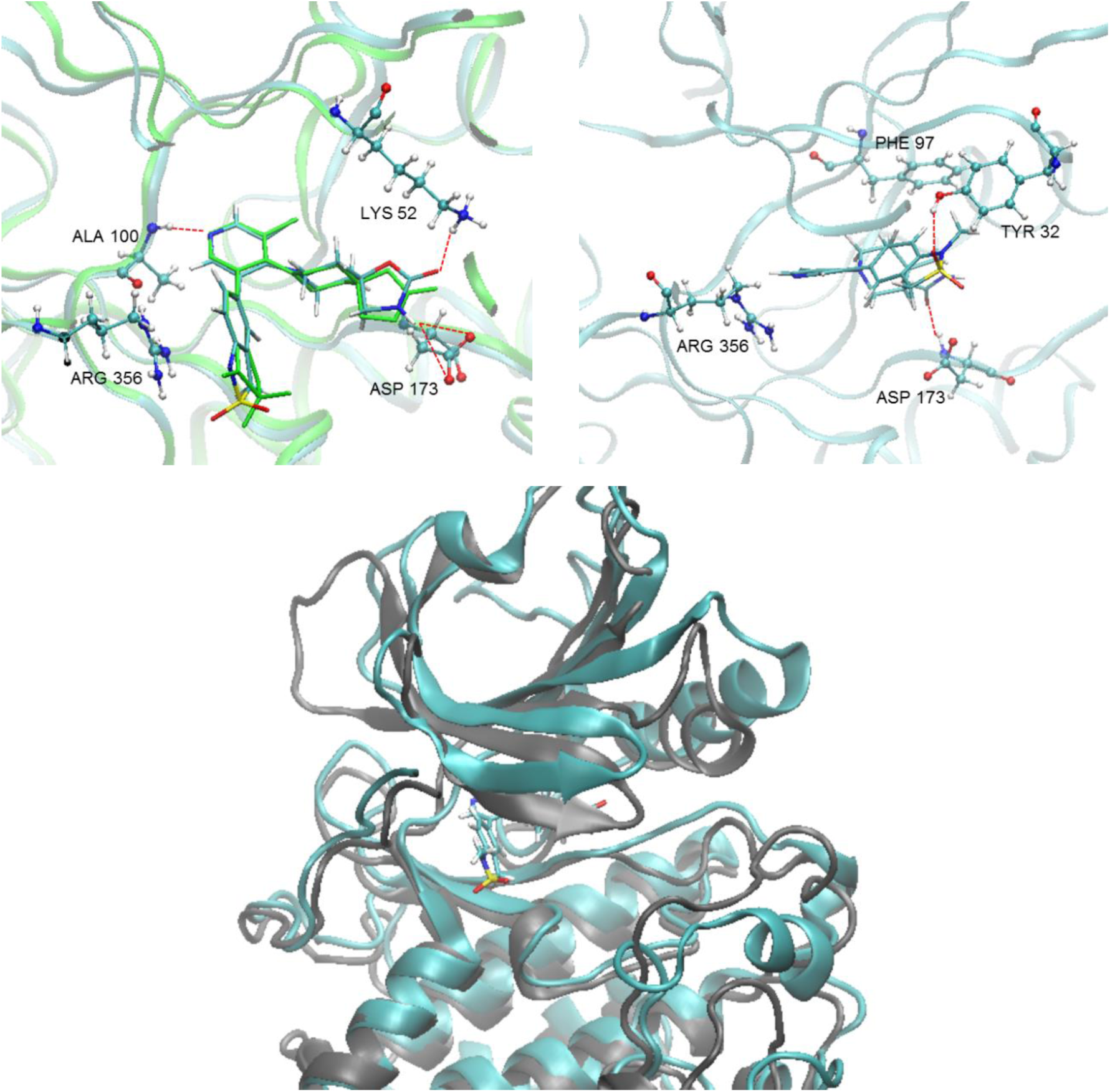
Top: Two binding modes observed in the complex of CDK8 and 5Y6 when CycC is absent, one at 0 ns (left) and the other at 200 ns (right). The crystal structure is colored green in the picture on the left side, aligned with the frame at 0 ns. Bottom: Superposition of the two frames with gray at 0 ns and cyan at 200 ns. Only the ligand at 0 ns is shown for clarity.

All the results as discussed above suggest that CycC stabilizes CDK8 by reducing the fluctuation of all its regions. The stability of CDK8 is important to ligand binding. MMPB/SA results show that in the presence of CycC ligands have stronger interactions with CDK8. Therefore, CycC promotes ligand binding as well. Furthermore, the activation of CDKs requires the binding of Cyclins, and phosphorylation of THR, SER, TYR on their activation loop [57, 58]. The binding of CycC changes the conformation of CDK8 dramatically [58, 28], and enables the ligand binding to the allosteric binding [25]. This agrees with our observation that when CycC is absent, the αC helix of CDK8 adopts an αC-out conformation, in which the GLU66 moves away from the DMG motif. By losing the H-bond from GLU66 and interactions from the entire αC helix, the allosteric binding site collapses, and this disables the binding of type II ligands.

### 3.3 Implications for Drug Design

In general, the major binding modes of the 10 CDK8 ligands obtained with our MD simulations are in good agreement with the crystal structures. In addition, our results provide insights into the driving forces of binding which are valuable for improvement of the existing ligands. For the DMG-in conformation and type I ligands, we observed significant decrease of flexibility at the region of the C-terminal tail. The fluctuation of the C-terminal tail is restricted by the strong cation-π interaction between ARG356 and type I ligands, while the flexibility of other protein regions is not influenced by ligand binding. A ligand which doesn’t have interaction with ARG356 may reduce the protein entropy penalty and thus increase affinity. This is supported by the recently discovered 4,5-dihydrothieno[3’,4’:3,4] benzo[1,2-d] isothiazole derivative which achieved a sub-nanomolar potency [26]. Based on a docking model of this ligand bound to CDK8, there is no contact between ARG356 and the ligand. The rigidity of type I ligands to the ATP binding site is also the key to achieve high affinity. All of the 5 type I ligands we studied have rigid scaffolds and are pre-organized to a V shape which fits snugly inside the pocket. However, 5Y7 has a relatively poor affinity due to its flexible pyrrolidine ring and the attached linear methoxyethyl group. Unlike other ligands, this flexible moiety of 5Y7 appears to be easily affected by the motion of the activation loop and could not gain stable contacts with its surrounding residues. Furthermore, since the driving force for the binding of type I ligands is dominated by vdW interactions, a ligand with more extensive contacts with the binding pocket residues such as PHE97, TRP105 and HIS106 may gain extra affinity.

For the DMG-out conformation and type II ligands, we observed significant decrease of flexibility of the αC helix due to the key H-bonding network involving GLU66 on the helix. Because of the importance of this H-bonding network to the binding of type II ligands, we cannot simply remove this interaction to improve affinity. Our MMPB/SA calculations overestimated the binding energies of type II ligands, suggesting that these ligands might pay high entropy penalty upon binding. Considering the flexibility of the type II ligands we studied, it is not surprising to see high entropy penalty. Therefore, to reduce the flexibility of type II ligands may be one feasible direction to ligand optimization. Meanwhile, interaction with the hinge region is another measure for improvement. Our MD results show that the hinge region acts as the pivot for the major protein motions. Contacts with this region would be subject to least influence from the protein motions once ligands are bound. The ligands we studied have no or merely intermittent interactions with the hinge region, so there is room for improvement.

The effect of mediating water molecules to the ligand binding is worth to be discussed as well. Mediating water molecules can be critical to ligand binding, but such information is usually difficult to be obtained from crystal structures. Because the identity of the mediating waters can change from ligand to ligand and many of the waters observed in an apo-structure are displaced by ligand binding, predicting the role of particular ordered water molecules in ligand binding remains challenging. Our MD results revealed some useful information about mediating water molecules in CDK8/CycC systems. Type I ligands forms relatively stable H-bonds with ASP173 through a mediating water molecule, with the occurrence percentage ranging from 22% to 37%. They also occasionally interact with GLU66 through a mediating water molecule, but this H-bond network appears unstable. Besides, water molecules also participate in the H-bonds between 5XG and TYR32, ASP103, ALA155, between 5Y7 and ASP103, HIS106, ARG356, between 5Y8 and ASP98, ASP103. The occurrence percentages of these H-bonds are between 22% and 52%. Among type II ligands, SKR1 is the only one which interacts with multiple residues including VAL27, TYR32, LYS52, ALA155, ASN156 and ARG356 through mediating waters. Other type II ligands interact with ASP98 and ALA100 at the hinge region either directly or through mediating waters. Some of the water-mediated H-bonds are quite stable and may have impacts on ligand binding. The detailed evaluation of the mediating water molecules is out of the scope of this paper.

Our MD results show that the “Crystal structure like” conformations are stable and prevailing, as evidenced by the fact that the predicted binding modes are generally in good agreement with those in crystal structures. The MD results also provide dynamics that is absent in crystal structures. Due to the protein motions, the interactions between residues and ligands could change along the MD time. Figure SI-4 records the H-bonds breaking and reforming in the course of 500 ns. Specifically, MD simulations offered us new conformations that are unavailable in crystal structures. We discovered a new binding mode of the αB helix to CycC on the complex of CDK8/CycC and 5Y6 after 300ns of MD run at 298 K. We found the frequent flipping of the pyrrolidine ring on 5Y7, and this motion appears to correlate to the motion of the activation loop. Furthermore, the MD results revealed the conformations of the activation loop missing from the crystal structures. The activation loop is highly flexible in nature, as shown in Figure 13. Our MD work thus provides the most wanted conformations of the activation loop for other computational research on CDK8. More importantly, despite of the high flexibility of the activation loop and its closeness to the binding sites, the binding modes of the ligands are mostly not affected by its motion. Such information is valuable to docking and free energy calculations.

**Figure 13.**
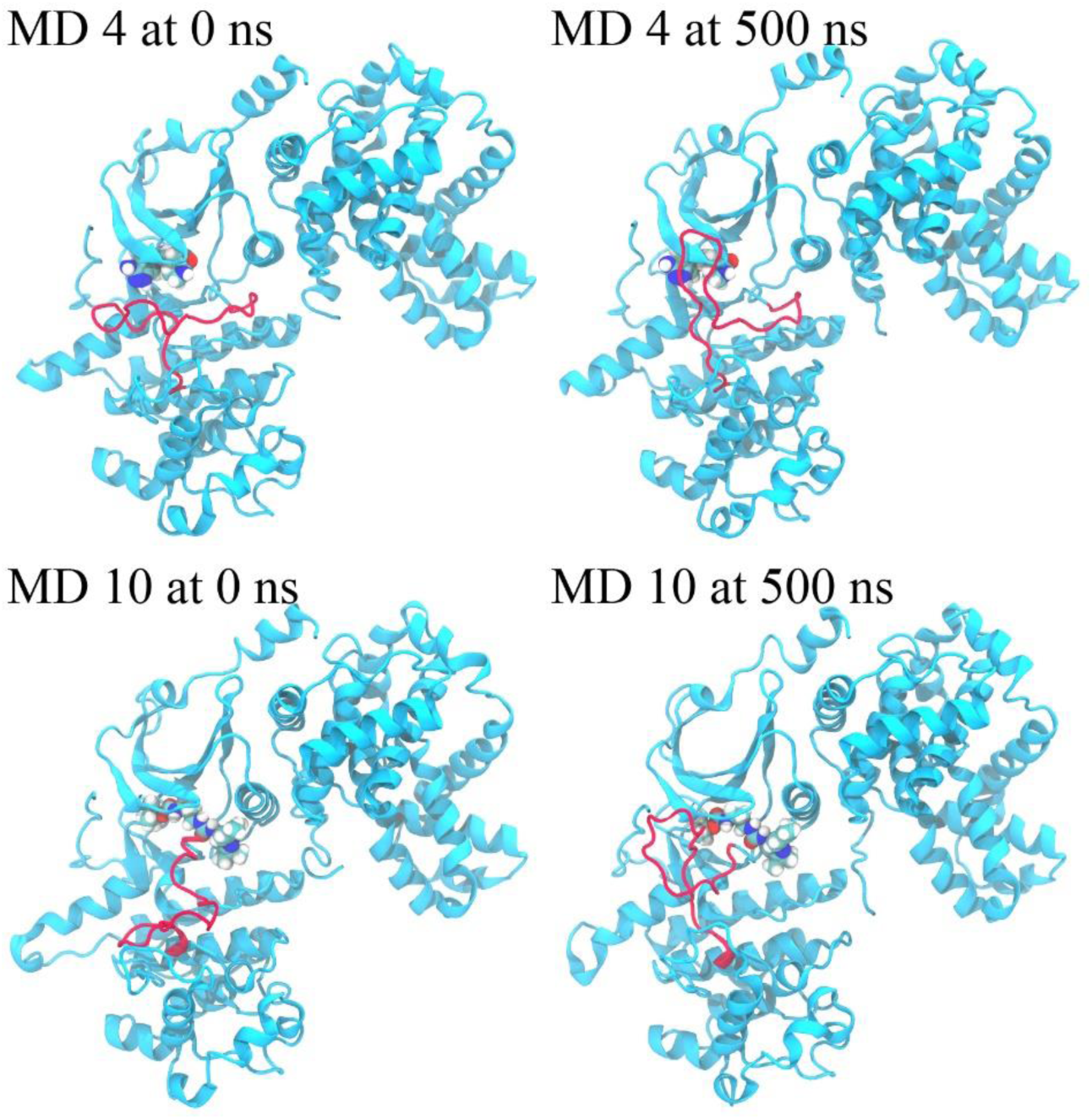
Highly flexible activation loop in both DMG-in (top) and DMG-out (bottom) conformations. Activation loop is colored red.

We calculated the total volumes of the ATP binding site and allosteric binding site for all the 13 systems. The average values and the deviations, together with a few representatives of the volume change along the MD time, are presented in Figure 14. Apparently the volumes are CDK8 conformation dependent. The apo protein in DMG-in conformation has smaller volume than the complexes, while the apo protein in DMG-out conformation has larger volume than the complexes. In general, the volumes of apo proteins have large fluctuations along the MD time, as indicated by the deviations, than those of complexes, presumably because the ligands interact with the protein and thus stabilize it.

**Figure 14.**
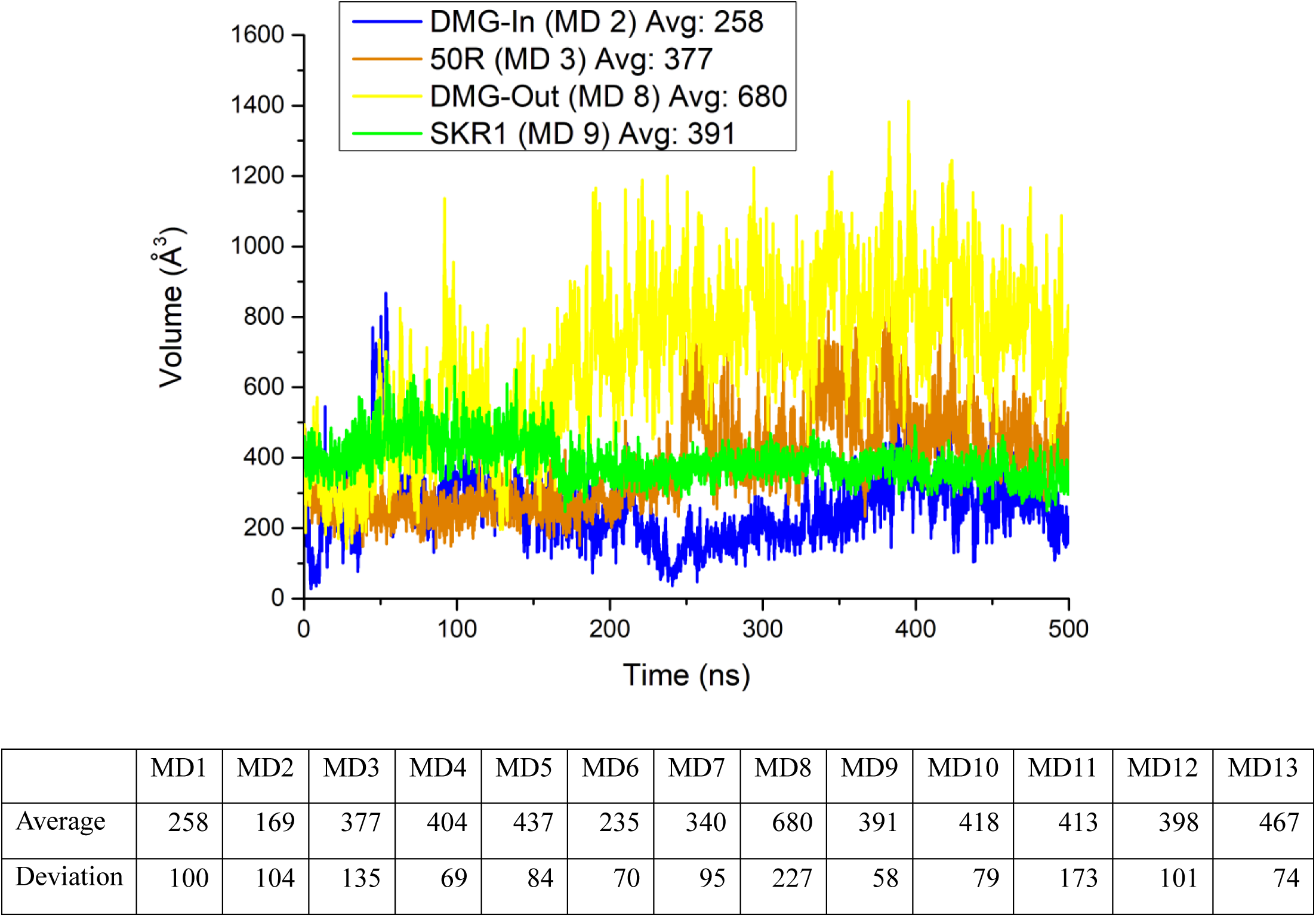
Volumes of the ATP binding site. Plot: Volume change along the MD time for four selected systems; Table: average volumes and deviations of all 13 MD systems.

The volume results suggest that the induced-fitting theory applies in the binding between type I ligands and the CDK8 DMG-in conformation. The ATP binding site starts with a small volume, and opens up to accommodate the ligands. The ligands push open the binding pocket, and the contacts are tight once they are bound. This is supported by the relationship between the average pocket volume and the experimental free energies for type I ligands, as in Figure 15. Interestingly, the pocket volumes have very high correlation with the binding affinities, with a R^2^ value of 0.946. Since the binding of type I ligands is mainly driven by van der Waals interactions, the larger volumes suggest more molecular contacts between the protein and the ligands, thus leading to higher affinities. An extrapolation of this relationship would suggest larger type I ligands for better affinities.

**Figure 15.**
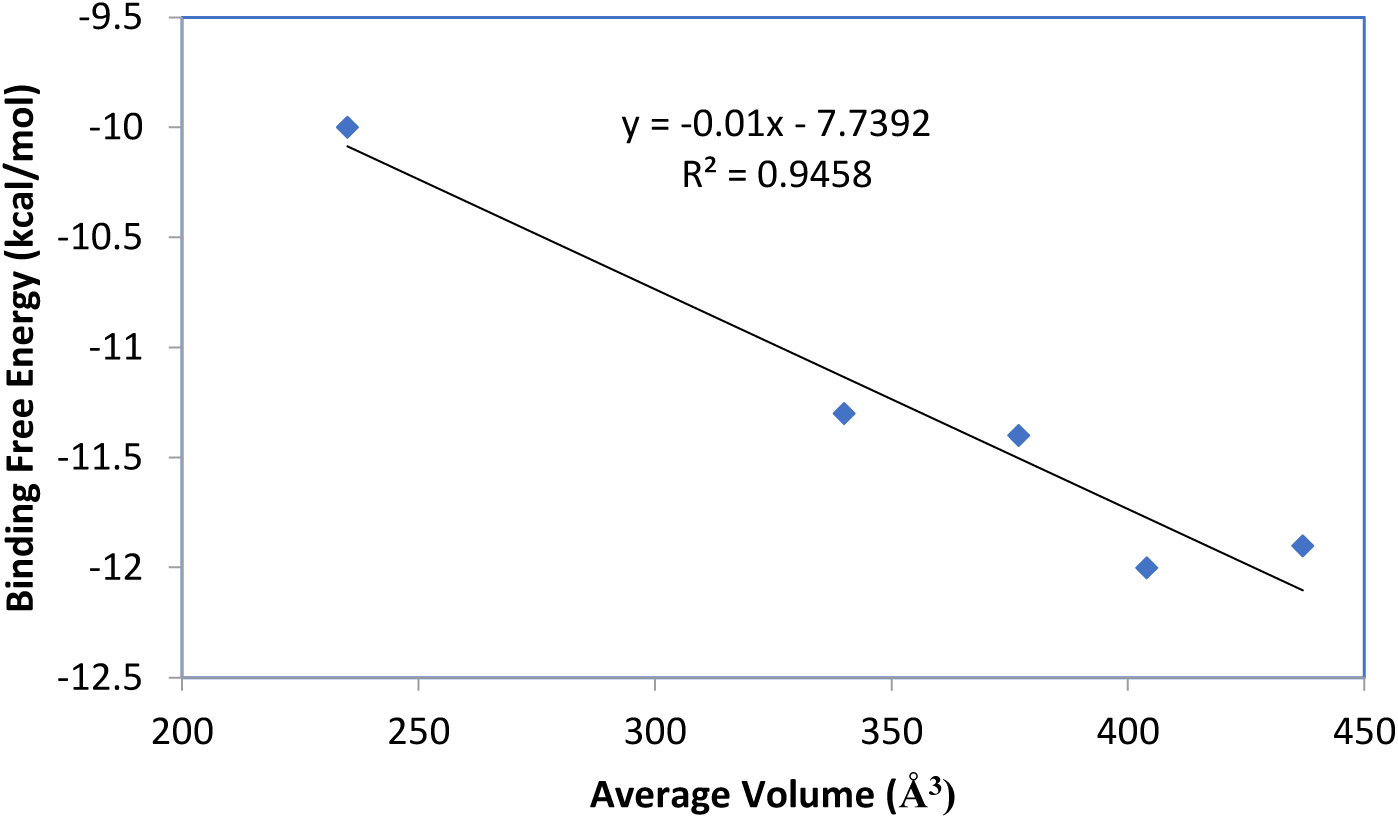
Relationship between the average volume of the ATP binding site and the experimental free energies for type I ligands.

The type II ligands seem to work in a different mechanism. These ligands have key interactions at the allosteric binding site and stretch into the ATP binding site to gain extra strength. The ATP binding site starts with a large volume. The ligands get into this region and “pull back” the surrounding residues to interact with them. The interactions between the protein and the ligands tighten the ATP binding site and stabilize it as well. SKR1 has the largest molecular size and is able to make most extensive contacts with the ATP binding site among the 5 type II ligands. It has the smallest pocket volume as well, indicating the tight contacts. The other type II ligands have smaller structures stretching into the ATP binding site, and the contacts are also smaller, as compared with SKR1. So the pocket tends to have larger volumes with these ligands. Since type II ligands have the major interactions at the allosteric site, it might not be necessary to fill up the ATP binding site to gain affinity. As aforementioned, the direction to improve type II ligands is to reduce their structural flexibility and to make contacts with the hinge region.

## 4. Conclusion

In this work we studied the dynamic behaviors and energy profiles with all-atom unbiased MD simulations for 13 CDK8/CycC systems, involving the DMG-in/out conformations, five type I ligands and five type II ligands. Our MD calculations provide interesting dynamics that is not available from crystal structures. The protein vibrates in the 500ns MD runs. All the large motions occurred roughly at the same regions in all 13 systems. The motions of the protein regions have high correlations. They move in concert and give five major protein motions. The hinge region between the N lobe and the C lobe acts as a pivot of the protein motions. Because the protein motions are related to the regions with large fluctuations, when these regions change along the simulation time the protein motions could also change. Due to the protein motions, some interactions between the protein and the ligands keep on and off along the MD time as well. People can thus investigate the binding mechanisms and find out alternative binding modes according to the stability and changes of the interactions from frames of MD runs. Different ligands cause fluctuations to different extents as indicated by RMSF values in some major regions such as the activation loop. However, the discrepancy in fluctuations didn’t seem to significantly affect the ligand binding modes. When compared with the motions of apo proteins, type I ligands remarkably reduce the motion of the C-terminal tail through the strong cation-π interaction between the ligands and ARG356 and type II ligands stabilize the αC helix by forming stable H-bonds with GLU66.

Some new conformations are discovered in the MD calculations. In addition to the conformations of the activation loop that are missing in the crystal structures, we found a new binding mode of the αB helix to CycC and the dynamics of the pyrrolidine ring on ligand 5Y7 which could be the reason for the relatively poor affinity of this ligand. All of the information would be valuable to the computational research on CDK8.

Our MD calculations confirm the importance of CycC to the stability of the CDK8 system as well as the ligand binding. CycC significantly reduces the motions of the αB helix, the αC helix and the activation loop. And due to the smaller motions of these regions ligands gain tighter contacts with CDK8.

The predicted binding modes of both type I and type II ligands resemble those in the crystal structures in general. Type I ligands have rigid molecular structures and type II ligands have more diversified and flexible structures. As a result, the MMPB/SA binding energies, which are lack of the entropy term, of type I ligands have relatively good correlation with the experimental free energies. The correlation for type II ligands is much poorer. Since type I ligands interact with ARG356 and restrict the motion of the C-terminal tail which can result in an entropy penalty to binding, to remove this interaction may help to improve the affinities of type I ligands. For type II ligands, to reduce their flexibility and increase the interactions with the hinge region may be the direction for improvement.

The calculations on the volumes of the ATP binding site for all the 13 systems reveal something of interest. The volumes in the DMG-in conformation have very high correlation with the binding affinities. The ATP binding site starts with a small volume, and opens up to accommodate the ligands. Thus, it appears that the induced-fitting theory applies in the binding between type I ligands and the CDK8 DMG-in conformation. This result also suggests that making type I ligands larger could obtain stronger affinity. For type II ligands, they have key interactions at the allosteric binding site. The ATP binding site of the DMG-out conformation starts with a large volume, and shrinks due to the contacts with type II ligands.

In summary, our MD simulations successfully revealed information on CDK8/CycC systems which is complementary to crystal structures. This information may be useful in the development of new CDK8 inhibitors and has implications in the field of drug discovery.

## Acknowledgements

We thank support from the US National Institute of Health (GM-109045), US National Science Foundation (MCB-1350401), and NSF national super computer centers (TG-CHE130009).

## Supplement Materials

**Figure SI-1.**
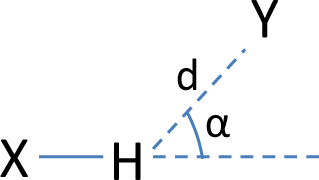
Definition of hydrogen bonds. X and Y stand for the donor and the acceptor respectively. d is the distance between the acceptor Y and the hydrogen, and α is the complimentary angle of X-H Y. A hydrogen bond is formed if d is smaller than 2.5 Å and α is smaller than 30°.

**Table SI-1.** MMPB/SA energy breakdowns for the binding interactions of 10 ligands with CDK8/CycC. Units: kcal/mol.

| Type I |  |  |  |  |  |
| --- | --- | --- | --- | --- | --- |
| MD Index | 3 | 4 | 5 | 6 | 7 |
| Ligand | 50R | 5XG | 5Y6 | 5Y7 | 5Y8 |
| PDB ID | 5CEI | 5FGK | 5HBE | 5HBH | 5HBJ |
| $\Delta E_{vdW}$ | -39.8±2.8 | -45.9±2.7 | -48.5±2.9 | -44.5±3.9 | -43.9±3.0 |
| $\Delta E_{Coulombic}$ | -13.0±4.6 | -28.7±6.4 | -24.6±7.0 | -90.8±9.2 | -26.0±5.5 |
| $\Delta E_{PB}$ | 31.2±3.6 | 47.2±5.7 | 48.0±6.6 | 117.7±8.6 | 41.4±4.9 |
| $\Delta E_{Non-Polar}$ | -3.6±0.1 | -4.2±0.1 | -4.3±0.1 | -4.8±0.2 | -4.2±0.1 |
| $\Delta E_{Gas}$ | -52.8±5.7 | -74.6±6.8 | -73.1±8.1 | -135.3±9.0 | -69.9±6.1 |
| $\Delta E_{Solv}$ | 27.6±3.5 | 43.1±5.6 | 43.7±6.6 | 112.9±8.6 | 37.3±4.9 |
| $\Delta E$ | -25.2±4.1 | -31.5±4.1 | -29.4±4.4 | -22.4±5.5 | -32.6±3.9 |
| $\Delta E_{Expt}$ | -11.4 | -12.0 | -11.9 | -10.0 | -11.3 |
| Type II |  |  |  |  |  |
| MD Index | 9 | 10 | 11 | 12 | 13 |
| Ligand | SKR1 | SKR2 | SKR5 | SKR10 | SKR11 |
| PDB ID | 4F6W | 4F7L | 4F6U | 4F7N | 4F7N |
| $\Delta E_{vdW}$ | -88.5±3.4 | -62.4±3.3 | -59.9±4.9 | -50.6±2.8 | -49.5±3.0 |
| $\Delta E_{Coulombic}$ | -20.4±4.1 | -20.0±4.8 | -28.3±7.6 | -20.9±4.6 | -19.0±6.0 |
| $\Delta E_{PB}$ | 66.4±4.7 | 49.9±5.0 | 52.6±9.0 | 40.4±5.0 | 38.3±5.4 |
| $\Delta E_{Non-Polar}$ | -7.9±0.1 | -5.8±0.1 | -5.5±0.1 | -4.8±0.1 | -5.0±0.1 |
| $\Delta E_{Gas}$ | -108.8±4.8 | -82.4±5.9 | -88.2±10.4 | -71.5±4.6 | -68.5±6.8 |
| $\Delta E_{Solv}$ | 58.5±4.7 | 37.0±5.0 | 47.1±9.0 | 35.6±5.0 | 33.3±5.4 |
| $\Delta E$ | -50.3±4.7 | -45.3±4.9 | -41.1±5.0 | -35.9±3.9 | -35.2±4.5 |
| $\Delta E_{Expt}$ | -10.2 | -10.7 | -8.4 | -8.0 | -9.7 |

**Figure SI-2.**
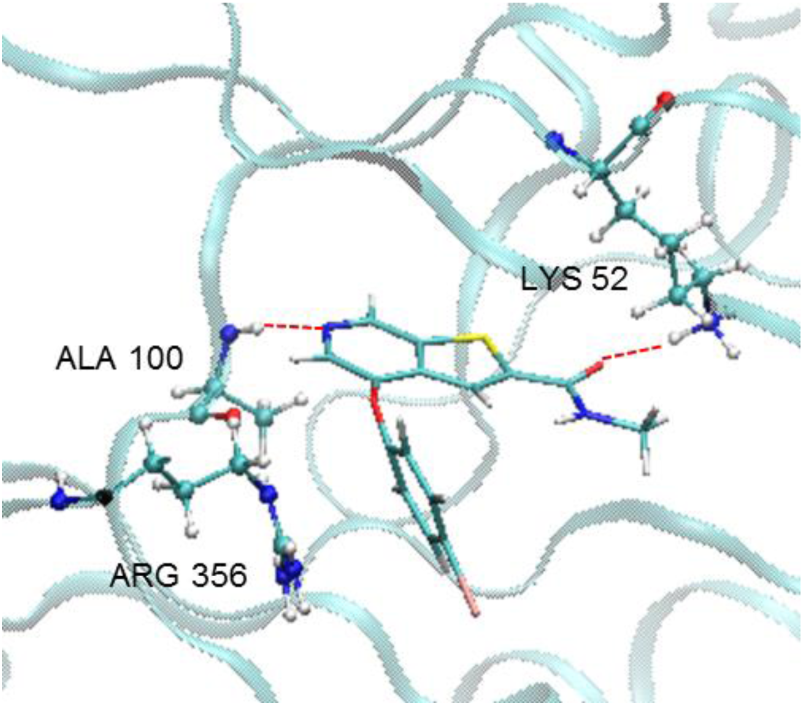
A typical binding mode of type I ligands with CDK8.

**Figure SI-3.**
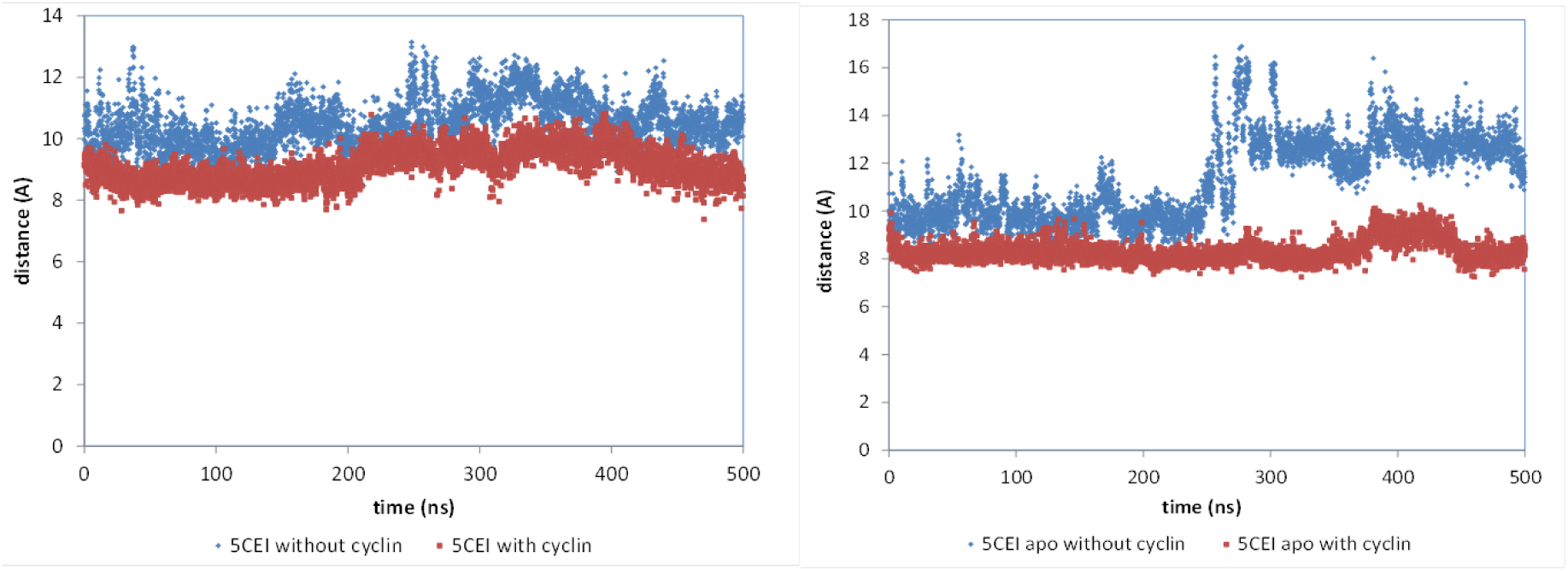
DMG-αC-helix distance changes with and without CycC in the CDK8 complex and apo protein.

**Figure SI-4.**
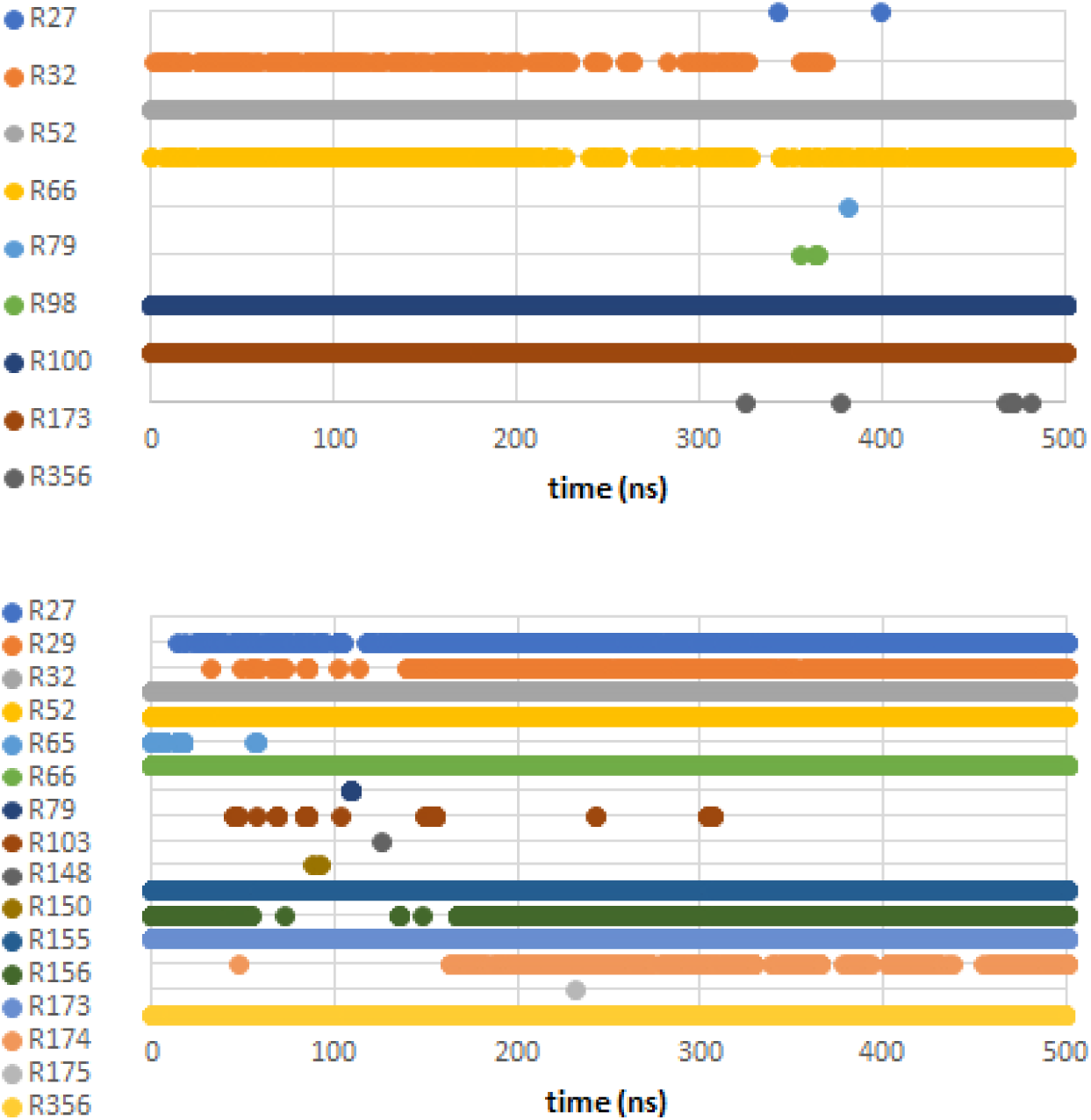
Breaking and reforming of hydrogen bonds between residues and ligands are observed along the MD time. Top: 50R; Bottom: SKR1.

